# A multiplexed in vivo approach to identify driver genes in small cell lung cancer

**DOI:** 10.1101/2022.03.28.485708

**Authors:** Myung Chang Lee, Hongchen Cai, Christopher W. Murray, Chuan Li, Yan Ting Shue, Laura Andrejka, Andy L. He, Alessandra Holzem, Alexandros P. Drainas, Julie H. Ko, Garry L. Coles, Christina Kong, Shirley Zhu, ChunFang Zhu, Jason Wang, Matt van de Rijn, Dmitri A. Petrov, Monte M. Winslow, Julien Sage

## Abstract

Small cell lung cancer (SCLC) is a highly lethal form of lung cancer. The high mutation burden in SCLC cells makes it challenging to predict key drivers of SCLC from genome sequencing data, thereby hindering the identification of possible therapeutic targets. Here we develop a quantitative multiplexed approach based on lentiviral barcoding with somatic CRISPR/Cas9-mediated genome editing to functionally investigate candidate regulators of tumor initiation and growth in genetically engineered mouse models of SCLC. Lentiviral vector-mediated SCLC initiation was greatly enhanced by naphthalene pre-treatment, enabling high multiplicity of tumor clones for analysis through high-throughput sequencing methods. Based on a meta-analysis across multiple human SCLC genomic datasets, we quantified the impact of inactivating 39 genes across many candidate pathways and captured both positive and detrimental effects on SCLC initiation and progression upon gene inactivation. This analysis and subsequent validation in human SCLC cells identified *TSC1* in the PI3K-AKT-mTOR pathway as a robust tumor suppressor in SCLC. This new approach should illuminate novel drivers of SCLC, facilitate the development of precision therapies for defined SCLC genotypes, and identify new therapeutic targets.

## INTRODUCTION

Small cell lung cancer (SCLC) comprises about 15 percent of all lung cancers and is one of the most aggressive forms of human cancer (Barnholtz-Sloan et al., 2004; Beasley et al., 2005; Travis et al., 2015). Mortality in SCLC remains high, with a median survival of 8-10 months, as SCLC tumors are highly metastatic and become rapidly resistant to therapeutic approaches (Reck et al., 2016). The malignancy of SCLC cells is at least in part encoded by the complexity of genomic alterations induced by cigarette smoking (George et al., 2015). A major goal in the field has been to identify genetic drivers of SCLC growth, with the intent of eventually mirroring the successes with targeted therapies achieved in lung adenocarcinoma (Howlader et al., 2020).

Inactivation of the *TP53* and *RB1* tumor suppressor genes is a near-universal event in SCLC. Other recurrent alterations in SCLC genomes include inactivation of tumor suppressors such as NOTCH family members or the MLL2 chromatin remodeler, and amplification of oncogenes such as MYC family transcription factors (Augert et al., 2017; George et al., 2015; Mollaoglu et al., 2017). Due to the high tumor mutation burden (TMB) in SCLC cells, however, distinguishing driver alterations from passengers remains challenging. For instance, *KMT2D* (encoding MLL2) is among the largest genes (>40kb) in the human genome; consequently, mutations within this gene have not been classified as putative drivers in the largest published genomic analyses of SCLC (George et al., 2015). However, functional analyses strongly suggest that MLL2 loss is an important driver of SCLC (Augert et al., 2017; Peifer et al., 2012).

Genetically engineered mouse models of cancer provide a platform with which candidate cancer driver alterations can be functionally interrogated in a relevant *in vivo* context. The development of SCLC mouse models that recapitulate the inactivation of *TP53* and *RB1* in human SCLC has enabled the investigation of the role of additional candidate drivers of SCLC (Calbo et al., 2011; Meuwissen et al., 2003; Schaffer et al., 2010). However, thus far, progress in functionally validating potential driver mutations has been slow, with only a limited number of genes being tested since the first mouse model of SCLC was developed close to 20 years ago (Ciampricotti et al., 2021; Coles et al., 2020; Cui et al., 2014; Denny et al., 2016; Jia et al., 2018; Kim et al., 2016; Mollaoglu et al., 2017; Schaffer et al., 2010).

Almost all genetically engineered mouse models of SCLC entail tumor initiation via the delivery of the Cre recombinase by an adenoviral vector (Ad-Cre) to delete conditional mutant alleles of *Rb1* and *Trp53* (Cui et al., 2014; Ferone et al., 2020; Meuwissen et al., 2003; Mollaoglu et al., 2017; Schaffer et al., 2010). Notably, adenoviral vectors do not integrate into the DNA of the transduced cells. While this may be beneficial in synchronizing the time of tumor initiation, it is unsuitable for experimental studies in which sustained transgene expression or genetic tagging of transduced cells is required. The recent development of Tuba-seq (<u>tu</u>mor <u>ba</u>rcoding with ultradeep barcode <u>seq</u>uencing) has enabled the functional investigation of pools of putative driver genes in a quantitative and scalable manner in mouse models of lung adenocarcinoma (Murray et al., 2019; Rogers et al., 2017). In this approach, each cell transduced by a lentiviral-Cre vector (Lenti-sgRNA/Cre) and its descendants are stably labeled with a clonal identifier in the form of a random DNA barcode (BC) as well as a vector-specific identifier to distinctly label each unique genetic perturbation (i.e., sgRNA-ID or sgID). Thus, the importance of each sgRNA-targeted gene during tumor initiation and growth can be studied quantitatively. Furthermore, the presence of sgID allows for simultaneous testing of multiple sgRNAs in one mouse with a pool of multiple lentiviral vectors. While the Tuba-seq pipeline is in theory generalizable to any *in vivo* model that is amenable to lentiviral transduction and relies upon a conditionally regulated tumorigenic program, it has not yet been applied to study the genetic underpinnings of SCLC development.

Here we present a barcoded Lenti-Cre-based mouse model of SCLC, which allows for tracking of SCLC tumor clones that develop entirely within the native environment. We show that pre-treatment with naphthalene is key to efficient initiation of SCLC using lentiviral vectors, which enables the analysis of many tumor genotypes with Tuba-seq. We quantitatively assessed the impact of over 30 genes on the initiation and growth of SCLC in a minimal number of mice. Our work validates the PI3K-AKT-mTOR pathway as an important driver of SCLC development and demonstrates a key role for TSC1 in this pathway as a potent tumor suppressor in SCLC.

## RESULTS

### Naphthalene treatment enhances the development of SCLC in mice

*Rb1^fl/fl^*;*Trp53^fl/fl^*;*Rbl2^fl/f^* (*RPR2*, or TKO, for triple knock-out) mice model the most prevalent subtype of human SCLC (SCLC-A, with high expression of ASCL1 (Rudin et al., 2019)). In this mouse model, tumor initiation is efficient and tumor progression relatively rapid (4-6 months) following intratracheal instillation with Ad-CMV-Cre (Gazdar et al., 2015; Schaffer et al., 2010). While *RPR2;R26^LSL-tdTomato^* (*RPR2T*) mice developed SCLC upon transduction with Lenti-Cre (HIV-PGK-Cre backbone) (**Figures 1A-C**), the overall tumor numbers were much lower than in *RPR2* mutant mice using Ad-CMV-Cre despite similar titers (**Table S1**) (Schaffer et al., 2010). Naphthalene is a compound that kills most club cells in the lung epithelium (Van Winkle et al., 1995). Based on a previous report using naphthalene prior to lentiviral transduction to generate lung tumors in mice (Xia et al., 2018), we injected *RPR2T* mice with either naphthalene or vehicle (corn oil) two days prior to intratracheal delivery of Lenti-Cre (**Figure 1A**). Naphthalene pre-treatment significantly increased both tumor number and burden in this context (**Figures 1B-C**). Importantly, Lenti-Cre-initiated tumors showed histopathological characteristics of SCLC-A tumors, including high expression of ASCL1 and the neuroendocrine marker UCHL1 (**Figures 1D** and **S1A**). Tumors initiated using a different Lenti-Cre vector backbone (FIV-CMV-Cre) showed similar SCLC-A histology (**Figure S1B**). In contrast to naphthalene pre-treatment, which increased tumor area also in the Ad-Cre model (**Figures S1C-D**), transduction with Ad-CMV-EGFP two days prior to Lenti-Cre transduction did not increase tumor number or area (**Figures S1E-F**), indicating that the pro-tumor effects of naphthalene are distinct from inflammatory responses upon adenoviral infection. Cell lines derived from *RPR2T* mutant tumors initiated by Ad-CMV-Cre or Lenti-Cre, with or without naphthalene treatment, formed floating clusters of cells in culture, similar to classical neuroendocrine SCLC cell lines (**Figure S2A**). Transcriptomic analysis confirmed expression of *Ascl1* and canonical neuroendocrine markers, with low levels of other transcription factors typical of other SCLC subtypes (Rudin et al., 2019) (**Figures 1F-G**).

**Figure 1:**
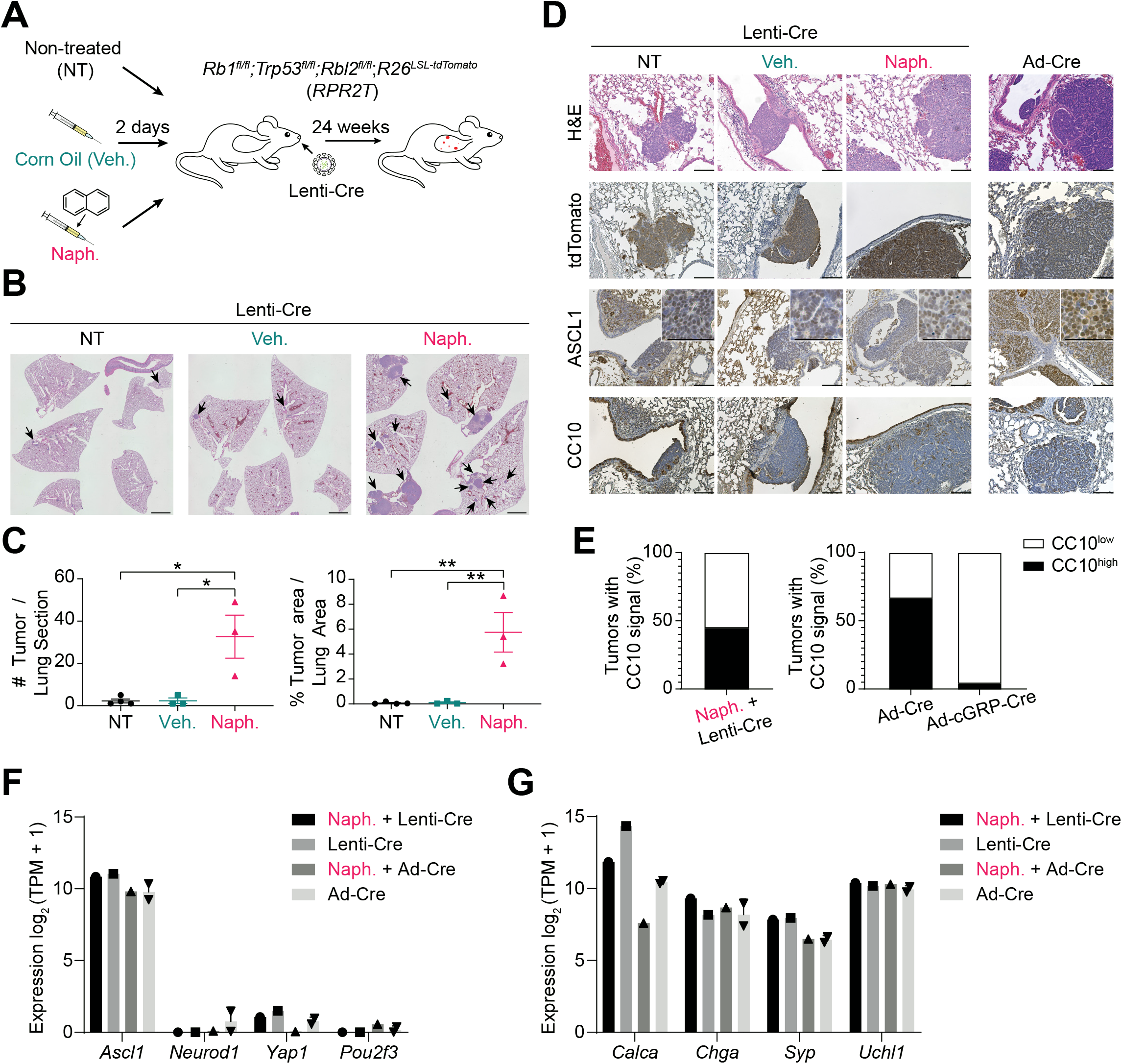
Naphthalene treatment enhances SCLC tumor development upon lentiviral Cre delivery. **A,** Workflow diagram for lentiviral Cre delivery (Lenti-Cre) to generate SCLC in mice (n=1 independent experiment). **B,** Representative hematoxylin and eosin (H&E) staining of lung sections (with some intestine in the middle panel) from mice transduced with Ad-CMV-Cre (Ad-Cre) or HIV-PGK-Cre (Lenti-Cre) alone (NT) or following corn oil (vehicle, veh.) or naphthalene (naph.) pre-treatment as in (A). Scale bar, 2 mm. **C,** Quantification of tumor burden and numbers from mice in (B) (n=3-4 mice per condition). P-values were calculated by one-way ANOVA with post-hoc Tukey test. *, p<0.05, **, p<0.01. **D,** Representative H&E and immunohistochemistry staining (IHC, brown signal) images of lung sections from mice transduced with HIV-PGK-Cre (Lenti-Cre) or Ad-CMV-Cre (Ad-Cre) as a control. Scale bar, 100 µm. Higher magnification images are shown in insets, where scale bar indicates 50 µm. **E.** Frequencies of CC10^high^ vs. CC10^low^ tumors quantified from images of lung sections from mice transduced with HIV-PGK-Cre (Lenti-Cre) as in (D) (n=2 mice). The analysis of tumors from mice infected with Ad-CMV-Cre (Ad-Cre) and Ad-cGRP-Cre are derived from data available in Yang *et al*., 2018. **F-G,** Bar graphs of RNA expression of selected genes (RNA-seq) in SCLC cell lines (Naph. + Lenti-Cre, n = 1; Lenti-Cre, n = 1; Naph. + Ad-CMV-Cre, n = 1; Ad-CMV-Cre, n = 2) (Lenti-Cre: HIV-PGK-Cre). (F) Genes representing the four major SCLC subtypes. (G) Common neuroendocrine markers. Data represented as mean ± s.e.m. (C) or mean ± s.d. (F-G).

SCLC tumors in the Ad-CMV-Cre *RPR2* model can be initiated from cGRP^+^ neuroendocrine lung epithelial cells (the minority of tumors) as well as from another, unknown non-neuroendocrine cell type(s) of origin (Yang et al., 2018). The presence of cells with non-neuroendocrine features (i.e., expressing the club cell marker CC10 or the NOTCH target HES1) within tumors initiated after naphthalene pre-treatment and Lenti-Cre transduction suggested that these tumors likely mostly originate from the same non-neuroendocrine cell type(s) as Ad-CMV-Cre SCLC tumors (Yang et al., 2018) (**Figures 1D-E** and **S2B-C**).

Thus, naphthalene pre-treatment enhances SCLC development in the *RPR2* model. Given the potential ease with which this Lenti-Cre platform could be used to inactivate genes of interest using sgRNAs and the CRISPR/Cas9 system, we next moved on to identifying potential regulators of SCLC etiology for further study.

### A meta-analysis reveals both known and novel putative genetic regulators of SCLC

To identify potential key drivers of SCLC pathogenesis, we performed a meta-analysis of 37 studies published prior to October 1^st^, 2021, on human SCLC (**Figure 2A**). We compiled data for genetic but also epigenetic and transcriptomic alterations in SCLC (**Table S2**). This analysis identified 3285 potential driver gene candidates that were profiled in at least 250 patients, had an alteration frequency of ≥ 3%, and coded for proteins with amino acid residue lengths of ≤ 2,000 (**Tables S3-4**). Whereas the relative rarity of SCLC tumor whole-genome/exome sequencing studies meant that many of the genes were examined in 400 patients or fewer, a number of the more highly profiled cancer-related genes benefitted from larger coverage, with the total patient numbers ranging from 500 to around 2000 (**Figure 2B**). As expected, in this analysis, inactivation of *TP53* and *RB1* ranked first and second, respectively, as the most frequently inactivated genes in SCLC (**Figure 2C**). While several top candidate genes identified in this meta-analysis, such as *COL11A1* and *XPC*, were previously identified to be recurrently mutated in SCLC or capable of initiating lung cancer upon inactivation (Hollander et al., 2005; Wagner et al., 2018), others, such as *HCN1*, *PCDH15*, and *ERICH3*, have been studied minimally in cancer contexts and represent novel tumor suppressor gene candidates in SCLC (**Figure 2C**). The 3285 candidates were enriched in signaling and cancer-related pathways (**Figure 2D**), including an enrichment in factors implicated in PI3K-AKT-mTOR signaling (**Figures 2D** and **S3A-B**, and **Table S5**). Furthermore, genes involved in DNA repair (Park et al., 2019), Notch signaling (George et al., 2015), the Wnt/Hippo-Merlin pathways (Wagner et al., 2018), and epigenetic and transcriptional regulation (Augert et al., 2017) showed high alteration rates (**Figure 2E**).

**Figure 2.**
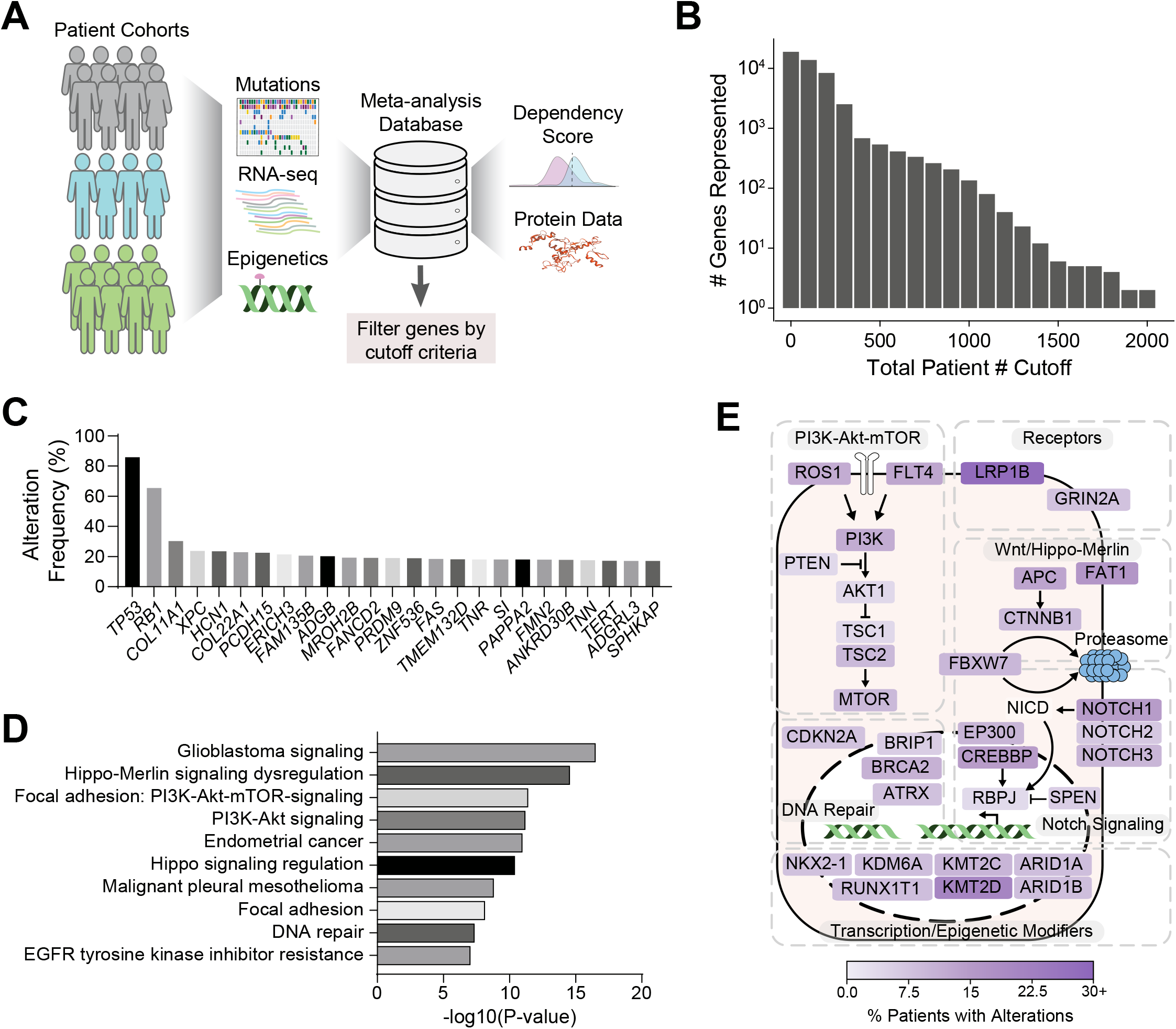
A meta-analysis of genetic studies identifies candidate drivers of SCLC development. **A,** Diagram of the meta-analysis workflow. **B,** Total number of genes represented when cutoff criteria are applied on total patient number (e.g., 2 genes were profiled in 2000+ patients, ∼100 genes were profiled in 1100+ patients). **C,** Alteration frequencies of top 25 gene candidates in all available patient data profiled. **D,** Top 10 enriched pathways for genes altered in ≥3% of SCLC patients, were profiled in at least 250 patients, and coded for protein with amino acid residue length of ≤2,000. Changing the gene cut-off criteria (e.g., removing amino acid residue length limits on protein products and keeping only genes that are expressed at ≥5 RPKM in human SCLC) did not strongly impact the pathway enrichments. WikiPathways was used for enrichment analysis, with the word “pathways” removed in the figure for space considerations. **E,** Diagram of selected SCLC driver candidates placed in signaling pathways based on (D). *RB1* and *TP53* were not represented to highlight other candidate drivers. Fill color indicates % of patients with alterations in that gene. P-value was determined through Bonferroni step-down correction on two-sided hypergeometric test (D).

Overall, this meta-analysis identified several candidate tumor suppressor genes and cancer pathways, most of which have not been functionally validated in SCLC (**Table S6**).

### Quantitative in vivo CRISPR screening to test tumor suppressive activity in SCLC

To investigate gene candidates in the pathways identified in the meta-analysis in an *in vivo* model of SCLC, we combined the Lenti-Cre/naphthalene platform with tumor barcoding with ultradeep barcode sequencing (Tuba-seq), an approach initially developed to uncover cancer drivers in mouse models of lung adenocarcinoma (Murray et al., 2019; Rogers et al., 2017). We transduced *RPR2T;H11^LSL-Cas9^* (*RPR2T;Cas9*) mice with four independent Lenti-sg*TSG*/Cre pools consisting of Lenti/Cre expressing inert sgRNAs (as controls) and Lenti-sgRNA/Cre vectors targeting putative tumor suppressor genes (TSGs), focusing on genes in key signaling pathways and novel targets from the meta-analysis (**Figure 3A**). Lungs were harvested at different time points for the different pools to identify possible optimal times for analysis, with no clear difference observed between the time points (see below).

**Figure 3:**
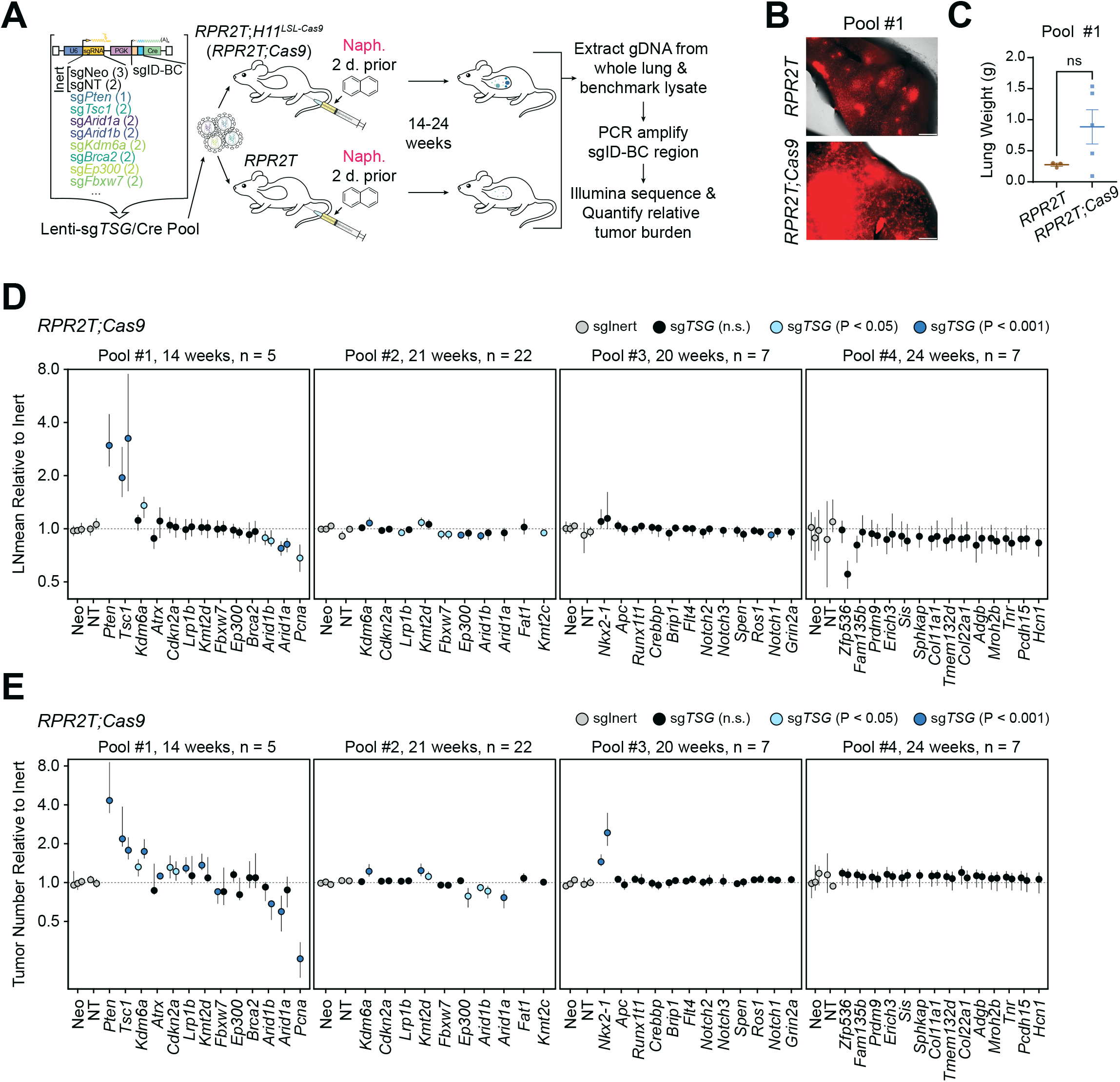
*In vivo* CRISPR screen uncovers both positive and negative effects of gene inactivation on SCLC growth and initiation. **A,** Diagram of the Tuba-seq workflow (n=4 independent experimental pools, n=3-22 mice per group). **B,** Lung fluorescence images from mice transduced with Pool #1. tdTomato fluorescence and brightfield images were merged. Scale bar, 1 mm. **C,** Lung weights of mice (n=3-5 per group) transduced with Pool #1 at the time of collection. **D,** Log-normal mean tumor size (normalized to tumors with sgInerts) for each putative tumor suppressor gene targeting sgRNA in *RPR2T;Cas9* mice. For each gene, each circle represents a unique sgRNA. P-values are indicated with a color code. **E,** Tumor numbers (normalized to tumors with sgInerts as well as tumors in *RPR2T* mice for Pools #1, 3-4 or *RPR2L* mice for Pool #2) for each putative tumor suppressor gene targeting sgRNA in *RPR2T;Cas9* mice. For each gene, each circle represents a unique sgRNA. P-values are indicated with a color code. The 95% confidence intervals were calculated by bootstrapping (D, E). p-values were determined through two-sided unpaired t-test (C) or bootstrapping followed by Benjamini-Hochberg correction (D, E). Data represented as mean ± s.e.m. (C) or mean ± 95% confidence interval (D-E). ns: not significant.

Both *RPR2T;Cas9* and control *RPR2T* mice showed robust tumor formation upon Cre delivery as evidenced by tdTomato fluorescence; *RPR2T;Cas9* mice showed a trend towards increased lung weights suggestive of increased tumor burden upon loss of tumor suppressor genes (**Figures 3B-C** and **S4A-B**). To determine gene inactivation effects on SCLC initiation and progression, we isolated genomic DNA from bulk tumor-bearing lungs, PCR-amplified and sequenced the sgID-BC region in the Lenti-sgRNA/Cre vector, and analyzed the data. A previous study using *RP;Pten^fl/fl^* conditional knockout mice showed that PTEN is a potent suppressor of SCLC development initiated by loss of p53 and RB (Cui et al., 2014). We found that inactivation of PTEN significantly increased tumor size and number in *RPR2T;Cas9* mice, indicative of a potent tumor suppressive role even with the additional inactivation of p130. Conversely, inactivation of the essential gene *Pcna* decreased tumor size and number, indicating that this pipeline has the ability to uncover both genotype-specific positive and negative effects on SCLC initiation and growth (**Figures 3D-E**) (Jaskulski et al., 1988). Distinct sgRNAs targeting the same gene consistently had similar effects in *RPR2T;Cas9* mice (**Figures 3D-E**). In contrast, sgRNAs targeting candidate cancer drivers had little to no effect on tumor growth in *RPR2T* mice lacking Cas9 (**Figures S4C-D**), as expected, indicating that sgID-BC enrichment and depletion recapitulate on-target gene inactivation activity. Quantifying tumor growth using alternative metrics showed similar results, with *Pten* and *Tsc1* inactivation increasing tumor burden and size and *Arid1a* and *Pcna* inactivation decreasing tumor burden in *RPR2T;Cas9* mice while sgRNAs having little to no effect in *RPR2T* mice (**Figures S5A-D**).

This Tuba-seq analysis identified several new genetic modifiers of SCLC growth (**Figures 3D-E**). First, *Tsc1* inactivation increased both tumor number and size, suggestive of a strong tumor suppressor role. Second, inactivation of *Arid1a* or *Arid1b* decreased tumor number and size in two independent pools, suggesting that ARID1A- and ARID1B-containing complexes normally promote rather than restrict the growth of SCLC in this genetic context. Third, inactivation of *Nkx2-1* led to increased tumor number but not size, suggesting a tumor suppressive role for this lung lineage transcription factor at the time of tumor initiation. Finally, *Kdm6a* inactivation increased tumor number in Pool #1, which was harvested at 14 weeks following tumor initiation, but showed a more modest effect in Pool #2 collected at 21 weeks following tumor initiation, suggesting that KDM6A may be a more potent tumor suppressor in early lesions in this model.

Taken together, these results indicate that adapting the Tuba-seq approach to autochthonous murine model of SCLC enables identification of oncogenic drivers and tumor suppressors, and greatly increases throughput of *in vivo* analyses.

### Single-guide validation confirms TSC1 as a tumor suppressor in mouse SCLC

Having observed frequent alterations in the PI3K-Akt-mTOR signaling pathway in our meta-analysis (**Figure 2E**) and identified *Tsc1* as a potent tumor suppressor gene in the *RPR2* model using Tuba-seq (**Figures 3D-E**), we further investigated the role of TSC1 in SCLC. In single-guide experiments, *RPR2T;Cas9* mice transduced with low titers of Lenti-sg*Tsc1*/Cre vectors (to ensure visualization of individual tumors) (**Table S1**) developed both SCLC tumors and NSCLC tumors (giant cell lung adenocarcinoma) (**Figures 4A-C**). Laser-capture microdissection followed by PCR amplification and sequencing of the sgID-BC in the Lenti-sgRNA/Cre vector showed that neighboring SCLC and NSCLC tumors arose from different clonal events (**Figures S6A-C**). While these *Rb/p53/p130/Tsc1* (*RPR2;Tsc1*) mutant tumors provide a new model for giant cell lung adenocarcinoma, we did not investigate their biology further. The neuroendocrine SCLC compartment in the *RPR2T;Cas9* mice with *Tsc1* inactivation showed a trend towards increased tumor number and area compared to control *RPR2T* mice at this time point (**Figures 4B-C** and **S7A**). A subset of the SCLC and NSCLC compartments had high levels of S6 phosphorylation compared to untransformed lung, suggesting elevated mTORC1 activity resulting from TSC1 inactivation (**Figure 4C**). S6 phosphorylation was also increased in *Tsc1*-KO SCLC cell lines compared to *Tsc1*-wt SCLC cell lines (**Figures 4D** and **S7B-C**).

**Figure 4:**
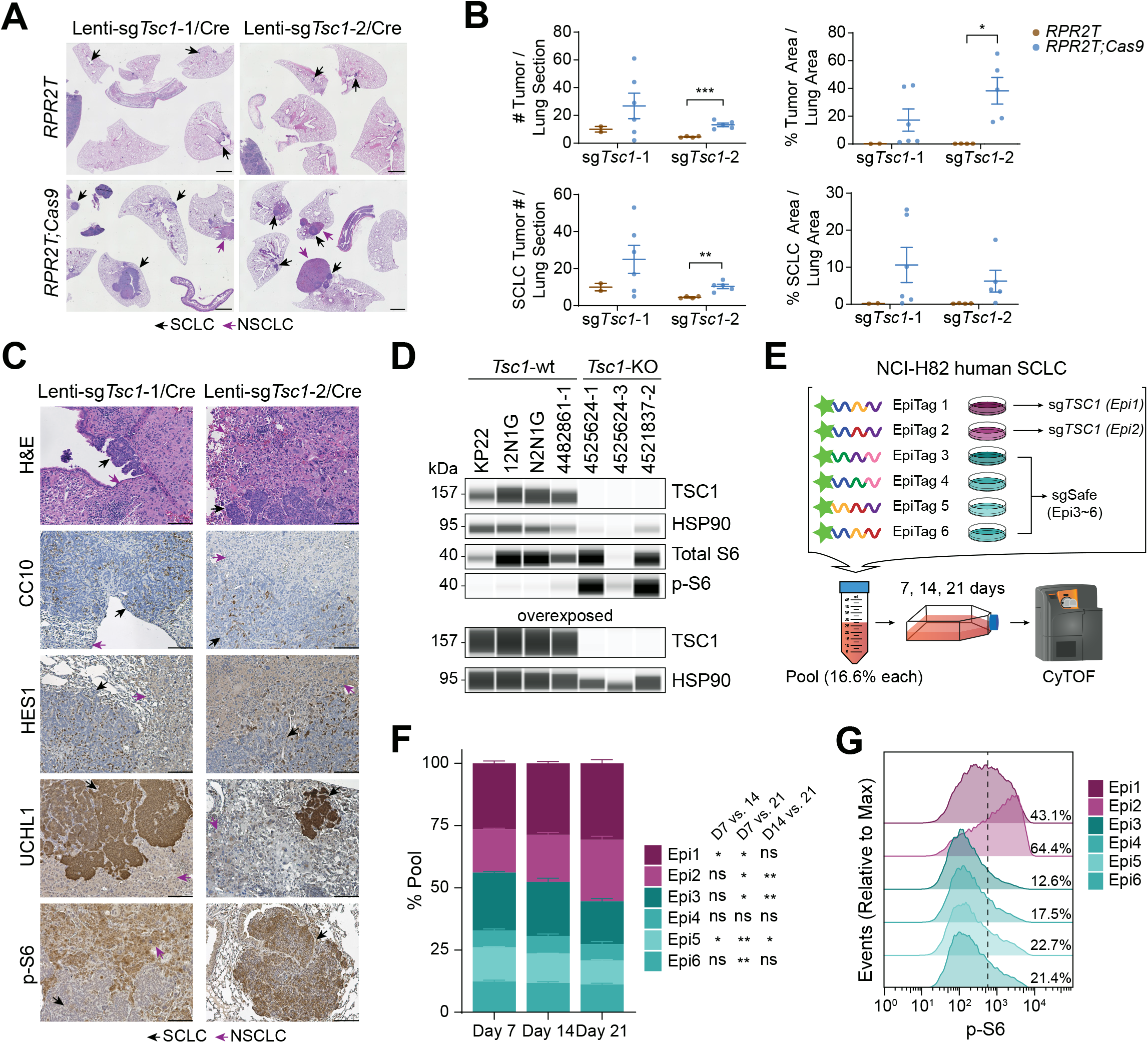
*TSC1* is a tumor suppressor in mouse and human SCLC. **A,** Representative H&E sections of lungs (and spleen and intestine) from *RPR2T* and *RPR2T;Cas9* mice transduced with Lenti-sg*Tsc1*/Cre sgRNA #1 (Lenti-sg*Tsc1*-1/Cre) or Lenti-sg*Tsc1*/Cre sgRNA #2 (Lenti-sg*Tsc1*-2/Cre) (n=2 independent experiments, 2-6 mice per group). Mice were collected 18 weeks after transduction. Scale bar, 2 mm. **B,** Quantification of tumor size and number in A. **C,** Representative H&E and immunohistochemistry staining (IHC, brown signal) images of lung sections from mice transduced with Lenti-sg*Tsc1*-1/Cre or Lenti-sg*Tsc1*-2/Cre. Scale bar, 100 µm. **D,** Immunoassay of TSC1, S6, and phosphorylated S6 (p-S6) in cell lines derived from *Tsc1*-wildtype (wt) and *Tsc1*-knockout (KO) mouse tumors. Overexposed image is shown to confirm the knockout. HSP90 was used as a loading control. **E,** Schematic of the pool competition assay (n=1 experiment with n=3 technical replicates) using Epitope-tagged (EpiTag) NCI-H82 cells. **F,** Stacked bar plot of percentage representation of epitope-labeled populations on Days 7, 14, and 21. Day 0 sample was unavailable. Statistical significance indicated next to epitope tags represent comparisons between Day 7 and 14, Day 7 and 21, and Day 14 and Day 21. **G,** Modal distribution of p-S6 signal across the different epitope-labeled populations. Percentage values represent the proportion of p-S6-high population. One representative experimental replicate from Day 21 is shown; all other replicates across days exhibit similar p-S6 signal distribution to what is shown. p-values were determined through two-sided unpaired t-test (B) or repeated measures two-way ANOVA with Geisser-Greenhouse correction followed by post-hoc Tukey test (F). ns, not significant; *, p<0.05; **, p<0.01; ***, p<0.001. Data represented as mean ± s.e.m. (B) or mean ± s.d. (F).

In the *RP* mouse model, Lenti-Cre initiates SCLC inefficiently even after naphthalene pre-treatment, with only 2-4 tumors visible at 42 weeks following transduction (**Figures S8A-B**). This low number of tumors and variable tumor development made quantification difficult and suggests that this mouse model may not be readily amenable to the Tuba-seq platform. *RP;R26^LSL-tdTomato^;H11^LSL-Cas9^ (RPT;Cas9)* mice also developed giant cell lung adenocarcinoma following transduction with Lenti-sg*Tsc1*/Cre vector but not the *RP;R26^LSL-tdTomato^ (RPT)* mice (**Figure S8C-D**), suggesting a broader role for *Tsc1* in regulating lung cancer development in different genetic contexts in mice.

The *RP* and *RPR2* models represent the SCLC-A subtype. In contrast, *Rb1^fl/fl^;Trp53^fl/fl^;H11^LSL-MycT58A^* (*RPM*) mice represent the Myc-overexpressing SCLC-N subtype (NEUROD1-high) (Mollaoglu et al., 2017). We generated *RP;R26^LSL-tdTomato/+^;H11^LSL-MycT58A/LSL-Cas9^* (*RPMT;Cas9*) mice and tested whether *Tsc1* acts as a tumor suppressor in this context also. As Lenti-Cre alone generate numerous tumors in the *RPM* model in only 8 weeks, we transduced *RPMT;Cas9* mice without naphthalene pre-treatment (**Figure S9A**). Compared to *RPMT;Cas9* mice transduced with Lenti-sg*Neo1*/Cre (Lenti-sgInert/Cre), mice transduced with Lenti-sg*Tsc1*/Cre had increased tumor number and area (**Figures S9B-C**). Of note, while *RPMT;Cas9* mice transduced with Lenti-sgInert/Cre generated NEUROD1-high SCLC, as expected, there were areas of NSCLC histology in *RPMT;Cas9* mice transduced with Lenti-sg*Tsc1*/Cre, consistent with our data in the *RPR2* and *RP* models (**Figures 4A** and **S9D**).

In all, our results indicate that TSC1 is a tumor suppressor in both ASCL1-high and NEUROD1-high subtypes of SCLC and that *Tsc1* inactivation is sufficient to induce NSCLC development in the *RPR2, RP,* and *RPM* genetic backgrounds.

### TSC1/TSC2 are tumor suppressors in human SCLC

Following validation of *Tsc1* as a tumor suppressor gene in mouse models of SCLC, we next sought to validate the tumor suppressive activity of TSC1 in human SCLC cells. We also investigated TSC2, the obligate partner of TSC1 within the tuberous sclerosis complex (Nellist et al., 1999). We generated populations of NCI-H82 SCLC cells independently expressing six unique combinations of short epitopes (epitope-combinatorial-tag, or Epi (Rovira-Clave et al., 2021)) and performed Cas9-RNA ribonucleoprotein nucleofection to generate *TSC1*-KO (in Epi1 cells), *TSC2*-KO (in Epi2 cells), and wild-type control cell lines (Epi3-6 cells, which received non-targeting Cas9-sgRNA ribonucleoprotein) (**Figures 4E** and **S10A**). We then pooled Epi1∼6-tagged NCI-H82 cells, cultured them for 21 days, and measured the relative change in epitope-tag representation using cytometry by time of flight (CyTOF) (**Figure 4E**). In this assay, *TSC1*-KO and *TSC2*-KO cells showed significantly increased expansion relative to the wild-type cell lines, making up the majority of the pool by Day 21 (**Figures 4F** and **S10B**). *TSC1*-KO and *TSC2*-KO cells also showed increased phosphorylated S6 compared to wild-type cell lines (**Figures 4G** and **S10C**). In contrast to the development of NSCLC upon loss of *Tsc1* at the time of initiation in the mouse models, inactivation of *TSC1* or *TSC2* in NCI-H82 cells did not lead to any obvious fate change towards a non-neuroendocrine fate, with no differences observed in markers indicative of epithelial-to-mesenchymal transition (EMT) and non-neuroendocrine differentiation (**Figure S10D**). These results validate *TSC1* and *TSC2* as tumor suppressor genes in human SCLC.

## DISCUSSION

In this study, we adapted a multiplexed and quantitative method to perform medium-throughput analysis of gene inactivation in mouse models of SCLC. Using this approach, we identified *TSC1* as a potent tumor suppressor gene in SCLC. The implementation of the Tuba-seq platform to mouse models of SCLC will greatly accelerate the functional analysis of candidate drivers of SCLC initiation and growth *in vivo*.

Lentiviral barcoding approaches have enabled breakthroughs in understanding tumor heterogeneity, clonal evolution, and multiplexed gene perturbation effects (Dixit et al., 2016; Nguyen et al., 2015; Wagenblast et al., 2015). In this study, we show that several advantages of lentiviral barcoding (e.g., multiplexed CRISPR screening and clonality analysis using barcoding approaches) can be captured in an autochthonous, *in vivo* context to study SCLC development. The application of Tuba-seq to mouse models of SCLC allowed us to investigate the gene perturbation effects at a much faster rate than previously capable. However, the lack of histological information (as exemplified with *Tsc1* loss and the appearance of a giant-cell carcinoma phenotype) and necessity for single-guide validation remain limitations of this approach. Nonetheless, the cost- and time-savings from using this approach to identify SCLC drivers cannot be overstated, and future approaches could take advantage of the lentiviral barcoding further, tracking metastasis drivers and dissecting tumor clonal evolution, for instance.

While the MSK-IMPACT panel remains a major resource for cancer genomics, including SCLC, its focus on readily-actionable cancer targets (341 to 468 genes) is a limitation (Bolton et al., 2020; Zehir et al., 2017). We sought to supplement and extend the currently available dataset in our meta-analysis from 37 different studies by adding more whole-genome/whole-exome sequencing studies in addition to studies profiling specific sets of genes. Because we aggregated our datasets for the sake of simplicity rather than keeping the individual patient IDs intact (i.e., keeping only the total number of patients with a given alteration in a gene), the cBioPortal remains a distinct resource for examining mutual exclusivity and co-occurrence patterns. Nonetheless, our meta-analysis database simplifies the search for novel cancer drivers by organizing aggregated patient alteration data alongside other useful metrics such as protein information for coding genes, RNA-seq expression datasets, and dependency scores from the Cancer Dependency Map (DepMap) project.

*TSC1*, alongside its complex partner *TSC2*, was first identified as a key gene whose mutation causes tuberous sclerosis complex (TSC) (Crino et al., 2006). TSC patients present with several clinical features, including skin lesions, hamartomas, and subependymal giant-cell tumor of the brain (Crino et al., 2006). The development of tumors in TSC patients, particularly giant-cell tumors, is thought to stem from TSC1-TSC2 complex’s role as a negative regulator of the mTOR pathway, which controls multiple pathways including cell growth (Brugarolas et al., 2004; Garami et al., 2003; Tapon et al., 2001). We observed mTORC1 activation and development of giant cell carcinoma of the lung upon *Tsc1* inactivation in our mouse models of SCLC. While only around 3-4% of SCLC patients have *TSC1* alterations, a substantial fraction of SCLC patients possesses PI3K-Akt-mTOR pathway alterations (e.g., 9-10% patients with *TSC2* alterations, 11-12% with *PTEN* alterations). Though cases of combined SCLC/giant cell carcinoma of the lung are rare in the clinic (Ebisu et al., 2018; Saito et al., 2017), our data suggest that these tumors may arise from dysregulation of the TSC1/TSC2-mTOR axis alongside RB/p53 loss-of-function.

Our data in mice and human cells show the strong tumor suppressive role of TSC1 in SCLC, as suggested by previous *in vitro* CRISPR/Cas9 knockout screens in mouse SCLC cell lines (Li et al., 2019). Several recent preclinical studies using SCLC models have also indicated that mTOR inhibition could be a viable strategy to treat SCLC patients, especially those resistant to chemotherapy. While rapamycin analogs (e.g., temsirolimus and RAD001), which preferentially inhibit mTORC1 rather than mTORC2, were met with little success in Phase II clinical trials (Pandya et al., 2007; Tarhini et al., 2010), ATP-competitive mTOR inhibitors (e.g., AZD-8055), which inhibit both mTORC1 and mTORC2, may be more promising. mTOR inhibition using AZD-8055 led to decreased tumor growth and sensitization to cisplatin/etoposide therapy in a subset of patient-derived xenograft models (Kern et al., 2020), and RICTOR amplification, which occurs in 10-15% of SCLC patients, was also shown to predict response to mTOR inhibitors in SCLC cell lines (Sakre et al., 2016). Furthermore, mTOR inhibition rescued the efficacy of Bcl-2 inhibition as well as WEE1 inhibition in *in vivo* models of SCLC (Gardner et al., 2014; Sen et al., 2017), leading to an ongoing Phase I/II clinical trial (NCT03366103). Taken together with our data, these results suggest that ATP-competitive mTOR inhibitors may produce therapeutic benefit in patients with alterations in the PI3K-AKT-TSC1/2-mTOR axis.

Increasing evidence from human tumor sections and mouse models indicates that intra-tumoral heterogeneity based on epigenetic/transcriptional program in SCLC cells plays a significant role in the growth of SCLC tumors and their response to treatment (Calbo et al., 2011; Gay et al., 2021; Lim et al., 2017; Mahadevan et al., 2021; Williamson et al., 2016). In contrast, our understanding of the genetic determinants of SCLC development has been hampered by limited tumor samples and the complex genome of these tumors. The development of medium-throughput pre-clinical approaches such as described here will contribute to a more rapid functional validation of genes and pathways relevant to SCLC in the clinic. A future goal of the field will be to explore how epigenetic and genetic mechanisms together mold SCLC development and response to various therapies to identify more personalized treatment strategies.

## MATERIALS AND METHODS

### Ethics statement

Mouse maintenance and experiments were conducted in accordance with practices prescribed by the NIH, the Institutional Animal Care and Use Committee (IACUC), and Association for Assessment and Accreditation of Laboratory Animal Care (AAALAC). The study protocol was approved by the Administrative Panel on Laboratory Animal Care (APLAC) at Stanford University (protocol #APLAC-13565).

### Mice and tumor initiation

*Rb1^fl/fl^;Trp53^fl/fl^;Rbl2^fl/fl^* (*RPR2*) mice has been described previously (Schaffer et al., 2010). *RPR2* mice were crossed with *Kras^LSL-G12D/+^;Trp53^fl/fl^*;*R26^LSL-tdTomato/LSL-tdTomato^;H11^LSL-Cas9/LSL-Cas9^* (*KPTC*) mice to generate *RPR2;R26^LSL-tdTomato/LSL-tdTomato^;H11^LSL-Cas9/LSL-Cas9^* (*RPR2T;Cas9*) and *RPR2;R26^LSL-tdTomato/LSL-tdTomato^* (*RPR2T*) mice. *Rb1^fl/fl^*;*Trp53^fl/fl^;R26^LSL-tdTomato/+^;H11^LSL-MycT58A/LSL-Cas9^* (*RPMT;Cas9*) mice were generated by crossing *Rb1^fl/fl^;Trp53^fl/fl^;H11^LSL-MycT58A/LSL-MycT58A^* (*RPM*) mice with *Rb1^fl/fl^;Trp53^fl/fl^*;*R26^LSL-tdTomato/LSL-tdTomato^;H11 ^LSL-Cas9/LSL-Cas9^* (*RPT;Cas9*) mice. 8- to 12-weeks-old mice were instilled with Lenti-sgRNA/Cre viruses via intratracheal delivery to generate lung tumors as previously described (DuPage et al., 2009). Viral titers used for experiments are detailed in **Table S1**. Ad5-CMV-Cre (Ad-Cre) and FIV-CMV-Cre (Lenti-Cre) were supplied by University of Iowa Viral Vector Core (VVC-U of Iowa-5 and VVC-U of Iowa-28).

Naphthalene (Sigma-Aldrich 184500) was dissolved into corn oil vehicle (Sigma-Aldrich C8267) at a concentration of 50 mg/mL and administered to mice via intraperitoneal (i.p.) injections at a dosage of 200 mg/kg.

### Cell line models

Human NCI-H82 SCLC cells (ATCC HTB-175™) and mouse SCLC cell lines (KP22, 12N1G, and N2N1G, described in (Denny et al., 2016)) were cultured in RPMI 1640 media (Corning 15-040-CV) supplemented with 10% bovine growth serum (BGS, Thermo Fisher Scientific SH3054103HI) and 1% Penicillin-Streptomycin-Glutamine (Gibco 10378-016). 293T cells used for lentiviral preparation were cultured in Dulbecco’s Modified Eagle Medium (DMEM) High-Glucose medium (Gibco 11965-118) supplemented with 10% fetal bovine serum. All cell lines were confirmed to be negative for mycoplasma (MycoAlert Detection Kit, Lonza LT07-418).

New mouse tumor-derived cell lines are described in **Table S7**. Briefly, tumor samples were microdissected and minced using a razor blade, digested with trypsin at 37°C for 5 minutes, quenched with RPMI media containing bovine growth serum (BGS), and centrifuged at 1000 RPM for 5 minutes to remove the supernatant. The pellet was resuspended in RPMI media, filtered through 40 µm membrane, and cultured at 37°C. Resulting tumor spheroids were checked for tdTomato fluorescence using Leica fluorescence microscope and imaged with LAS X software (v3.7.1, Leica Microsystems, Wetzlar, Germany). *In vitro* fluorescence images were pseudo-colored using Fiji (v1.53f51) (Schindelin et al., 2012).

### Ribonucleoprotein nucleofection with Cas9

Control and targeting sgRNAs were generated as previously described (Rovira-Clave et al., 2021). Briefly, for each region of interest, three sgRNAs were designed to hybridize approximately 150 bases apart, and 100 pmol of each sgRNA was resuspended in Tris-EDTA (Synthego) and mixed at a 1:1:1 ratio. In a 96-well v-bottom plate, 3 µL of the sgRNA (total 300 pmol) was added to 12 µL of SE buffer (Lonza V4XC-1032). In another well, 0.5 µL of Alt-R® S.p. Cas9 (Integrated DNA Technologies 1081059) was added to 10 µL of SE buffer, and the Cas9 mix was added to the sgRNA solution, mixed thoroughly, and incubated at 37°C for 15 minutes to form the RNP solution. 1 x 10^6^ NCI-H82 cells were resuspended in 5 µL of SE Buffer, and cells were nucleofected with Lonza 4D-Nucleofector™ X Unit (Lonza AAF-1002X) with the EN150 program immediately following addition of the RNP solution. Following the nucleofection, warm RPMI media was added to the cells, and cells were incubated at 37°C for 15 minutes then transferred to a 24-well plate.

### Cell preparation for CyTOF

Frozen cell lines in RPMI media supplemented with 10% BGS and 10% DMSO (Fisher Scientific BP231) were thawed, and 3 million cells per sample were washed once with PBS. The cells were then fixed with 1.6% formaldehyde at room temperature for 20 minutes. We are using palladium (Pd) barcoding to pool up to 20 different samples and reduce tube-to-tube variability. Therefore, cells were washed twice with PBS before permeabilization with PBS and 0.02% Saponin (Sigma-Aldrich 84510) at 4°C. 11 µL of Pd barcode was added to 1 mL of PBS and 0.02% Saponin, of which 900 µL were used to resuspend each sample. This mix was incubated at room temperature for 15 min, washed three times with Cell Staining Media (CSM, PBS with 0.5% BSA (Thermo Fisher Scientific B14), 0.02% NaN_3_ (Fisher Scientific MP210289110)), and then pooled into a single tube for staining with metal-labeled antibodies (**Table S8**) for 1 hour at room temperature. Antibodies against Flag, mCherry, GFP, VSV, NWS, Prot C, Ha, AU1, Synaptophysin, Vimentin, S6, GMNN, EZH2, pYAP, CDT1, pI3K, HES1 were conjugated with MAXPAR X8 Multimetal Labeling Kit (Fluidigm 201300) according to the manufacturer’s protocol. Antibodies were diluted to 0.2 µg/µL and titered. Cells were stained with a range of 1:100 to 1:200 with each of the different antibodies in a staining volume of 150 µl (∼ 3 × 10^6^ cells/mL). After antibody staining, the cells were washed twice with CSM and then incubated overnight at 4°C with an iridium-containing intercalator (Fluidigm 201192B) in PBS with 1.6% formaldehyde. The cells were then washed twice with water, diluted with water and 10 µL/ml EU Four Element Calibration Beads (Fluidigm 201078) to ∼ 10^6^ cells/mL, and filtered through a 70-μm membrane (Falcon 352350) just before analysis by mass cytometry.

### Lentiviral vector generation and titering

Lentiviral vectors containing individual sgRNAs, barcode sequences, and Cre recombinase were generated as previously described (Rogers et al., 2017). Briefly, sgRNA sequences were picked based on an aggregated score from top hits on Desktop Genetics (formerly www.deskgen.com) and GPP sgRNA Designer offered by the Broad Institute (https://portals.broadinstitute.org/gpp/public/analysis-tools/sgrna-design). Several of the sgRNAs used have been validated in previous studies (Cai et al., 2021; Rogers et al., 2017). Detailed sgRNA and barcoding primer sequence information can be found in **Table S9**. Lenti-sgRNA/Cre plasmids were barcoded individually with an 8-nucleotide ID specific to each sgRNA (sgID) and 20-nucleotide random barcode sequence (BC), and each plasmid was packaged separately in 293T cells via co-transfection with polyethylenimine alongside pCMV-VSV-G (Addgene #8454) envelope plasmid and pCMV-dR8.2 dvpr (Addgene #8455) packaging plasmid (Stewart et al., 2003). Sodium butyrate (Sigma-Aldrich B5587-5G) was added eight hours following the transfection to increase viral titer. Viral supernatant was collected at 48 and 60 hour time points following transfection, filtered using 0.45 µm PES syringe filter (Millipore SLHP033RB), concentrated via ultracentrifugation at 25,000 RPM for 90 minutes at 4°C, resuspended in PBS overnight, and titered using LSL-YFP mouse embryonic fibroblasts (MEFs) as previously described (Rogers et al., 2017).

### Histology, immunohistochemistry, and immunofluorescence

Mouse tissues were dissected from animals immediately following euthanasia. Lungs were inflated with 10% neutral buffered formalin (NBF), and all tissues were fixed in 10% NBF overnight following a brief rinse in PBS. Tissues were transferred to 70% ethanol prior to paraffin embedding and processing.

Prior to immunohistochemistry, paraffin sections were rehydrated by 5-minute serial immersion in Histo-Clear, 100% ethanol, 95% ethanol, 70% ethanol, and water. Antigen retrieval was performed by immersing rehydrated slides in citrate-based antigen unmasking solution (H-3300, Vector Laboratories) at boiling temperature for 15 minutes. To block endogenous peroxidase activity, the slides were then incubated in 3% hydrogen peroxide for 1 hour. Slides were washed in PBS-T (PBS + 0.1% Tween-20), blocked using blocking buffer (5% horse serum in PBS-T) for 1 hour at room temperature, and incubated with primary antibodies at 4°C overnight. Following incubation, slides were washed in PBS-T, incubated with the secondary antibody for 1 hour at room temperature, and developed using DAB reagent following another series of PBS-T washes. For HES1, ImmPRESS® Excel Amplified Polymer Staining Kit, Anti-Rabbit IgG, Peroxidase (Vector Laboratories MP-7601) or TSA Plus Fluorescein kit (Akoya Biosciences NEL741001KT) were used to amplify signal for immunohistochemistry or immunofluorescence, respectively. Following development, slides were counterstained using hematoxylin, dehydrated by 5-minute serial immersions in 70% ethanol, 100% ethanol and xylene, and mounted with mounting media.

For immunofluorescence, all the same steps as immunohistochemistry were followed until the secondary antibody step; slides were incubated with fluorescent secondary antibodies diluted in blocking buffer (5% horse serum in PBS-T) for 1 hour, washed in PBS-T, and stained with 0.6 nM DAPI in PBS for 10 minutes at room temperature. Slides were mounted with Fluoromount-G (SouthernBiotech 0100-01) and stored in 4°C overnight or -20°C for a few days before visualization.

The following antibodies were used for immunohistochemistry and immunofluorescence: ImmPRESS HRP Horse anti-Rabbit IgG (Vector Laboratories MP-7801-15), ImmPRESS HRP Horse anti-Mouse IgG (Vector Laboratories MP-7802-15), Alexa Fluor 594 Donkey anti-Goat IgG (H+L) Cross-Adsorbed Secondary Antibody (Invitrogen A11058), Alexa Fluor 488 Donkey anti-Rabbit IgG (H+L) Highly Cross-Adsorbed Secondary Antibody (Invitrogen A-21206), anti-RFP (for immunohistochemistry, Rockland 600-401-379, 1:200), anti-RFP (for immunofluorescence, MyBioSource MBS448122, 1:200), anti-CC10 (E-11, Santa Cruz sc-365992, 1:200), anti-HES1 (D6P2U, CST 11988S, 1:200), anti-UCHL1 (Sigma-Aldrich HPA005993, 1:2,500), anti-phospho-S6 (Ser235/236, CST 2211, 1:200), and anti-MASH1 (BD Biosciences 556604, 1:200).

### Quantitative immunoassay analysis

Cells were lysed in RIPA buffer (50 mM Tris-HCl pH 7.5, 1% NP40, 2 mM EDTA, 100 mM NaCl) supplemented with cOmplete™ ULTRA Protease Inhibitor Cocktail tablets (Roche 5892970001) and PhosSTOP™ phosphatase inhibitor tablets (Roche 4906845001). Pierce BCA Protein Assay Kit was used to quantify total protein concentration (Thermo Fisher Scientific 23227). For quantitative immunoassays, Simple Western™ assay was performed on the Wes™ system (ProteinSimple) according to manufacturer’s protocol, with 1 µg of protein loaded per lane. Primary antibodies against the protein of interest and a loading control protein were run simultaneously in each lane. Compass for SW (v5.0.1, SimpleWestern) software was used for protein quantification and size determination. The authors note that the protein sizes tend to run larger on the Wes system compared to traditional immunoblotting. The following antibodies were used: anti-HSP90 (CST 4877, 1:4,000), anti-TSC1 (D43E2, CST 6935, 1:200), anti-TSC2 (D93F12, CST 4308), anti-S6 (CST 2217, 1:200), and anti-phospho-S6 (Ser235/236, CST 2211, 1:200).

### Literature meta-analysis

Primary studies containing SCLC-specific alteration data were collated according to data availability and alteration profiling method. The patient and alteration counts for each gene were collected from data tables associated with the study where available. For studies without such data tables, the counts were determined by manual annotation of the OncoPrint figures. To determine the number of patients with alteration(s) as well as the total number of patients profiled for each gene, each of the two numbers were summed across all studies, and these two numbers were used to estimate the proportion of SCLC patients possessing an alteration for a given gene. The protein name and amino acid length associated with each gene were obtained through a UniProt DB query, and genes associated with multiple protein notations were each annotated with the overlapping UniProtKB entries (e.g. OR2A1, OR2A42 were both annotated with the UniProtKB entry for Olfactory receptor 2A1/2A42). To examine the expression level for each gene, we collated the data from both human SCLC samples as well as mouse models of SCLC (George et al., 2015; Yang et al., 2018). Percentile values for gene expression were determined from each dataset independently. To include a functional output from available *in vitro* experiment data, we also collected gene dependency scores from the CRISPR knockout screens (Achilles) conducted through the DepMap project (Dempster et al., 2019; Meyers et al., 2017). Cell lines were grouped according to cancer type (SCLC, NSCLC, and all cancers other than SCLC), and the median dependency score was calculated for each gene. AACR GENIE data was obtained from Synapse Platform with written permission (AACR Project GENIE Consortium, 2017).

### Preparation of Tuba-seq sgID-BC amplicon libraries

Genomic DNA was extracted from tumor-bearing mouse lungs following the addition of three benchmark control cell lines (1 x 10^5^ cells per control) as previously described (Rogers et al., 2017). Briefly, the lungs were homogenized and lysed with overnight protease K digestion, and genomic DNA was extracted from the lysate using phenol-chloroform and ethanol precipitation methods. Libraries were prepared by single-step PCR amplification of the sgID-BC region from 32 µg of gDNA per mouse split across eight 100 µL reactions with NEBNext Ultra II Q5 Master Mix (M0544L). Dual index primer pairs with unique i5 and i7 indices were used. PCR products were purified using Sera-Mag Select beads (GE Healthcare Life Sciences 29343052) and assessed for quality with Agilent High Sensitivity DNA kit (Agilent Technologies 5067-4626) on the Agilent 2100 Bioanalyzer (Agilent Technologies G2939BA). Purified libraries from each mouse were pooled at equal ratios based on lung weight to ensure even sequencing depth per cell, purified once more with Sera-Mag Select beads to remove excess free primers, and sequenced on the Illumina HiSeq 2500 or NextSeq 550 platform (Admera Health Biopharma Services).

### Tuba-seq analysis

We identified the target gene and random barcodes from the sgID-BC region for each tumor cell as previously described (Li et al., 2021). The absolute cell number in each tumor was calculated by normalizing the sgID-BC read number by that of the three benchmark control cell lines. We focused on tumors with at least 200 cells and calculated the LNmean (maximum likelihood estimator for mean tumor size assuming log-normal distribution) and tumor number for tumors carrying each target gene deletion. The LNmean and tumor number were normalized to that of Inert tumors to represent the relative growth advantage after inactivating these genes.

### Laser capture microdissection and tumor clonality analysis

7 µM sections from formalin-fixed, paraffin-embedded (FFPE) tissue blocks were cut and mounted on PEN membrane slides (Thermo Fisher LCM0522). Slide was dissected immediately after staining on an Arcturus XT LCM System (Thermo Fisher A26818). The cells in different regions were separated and adhered to CapSure HS LCM Caps (Thermo Fisher LCM0215). Genomic DNA was isolated from these different caps using PicoPure DNA Extraction kit (Thermo Fisher KIT0103). 50 µL lysis buffer with proteinase K were added into each sample and incubated at 65°C overnight. After inactivating proteinase K at 95°C for 10 mins, the genomic DNA was cleaned up with AMPure XP beads at 3:1 ratio (Beckman Coulter A63880) and eluted in the 10 mM Tris-HCl (pH 8.0). The DNA concentration was measured by Qubit® dsDNA HS Assay Kit (Thermo Fisher Q32851).

sgID-BC amplicon libraries were prepared from genomic DNAs by single-step PCR amplification with NEBNext Ultra II Q5 Master Mix (M0544L) using dual indexing primer pairs. PCR products were purified first using MinElute PCR Purification Kit (Qiagen 28006), then with Agencourt AMPure XP beads (Beckman Coulter A63881) and assessed for quality with Agilent High Sensitivity DNA kit on the Agilent 2100 Bioanalyzer prior to sequencing on the Illumina MiSeq Nano platform (Admera Health Biopharma Services).

### RNA-seq analysis

Tumor-derived cell lines were snap frozen and submitted for RNA analysis. Total RNA isolation, polyA selection, quality control, library preparation, and sequencing were performed by Azenta using Illumina HiSeq platform (2×150 bp, ∼350M paired-end reads). Transcript quantification for the RNA-seq data was conducted with Salmon (v0.12.0) (Patro et al., 2017) with mouse genome version GRCm38. DESeq2 (v1.34.0) was used to calculate differential expression across the mouse tumor-derived cell lines (Love et al., 2014).

### Statistical analysis

ClueGo plugin (v2.5.6) running on Cytoscape (v3.8.0) (Bindea et al., 2009; Shannon et al., 2003) was used to determine pathways enriched for 3285 genes that had been profiled for ≥ 250 patients, having an alteration proportion of ≥0.03, and coding for proteins ≤2000 amino acid residues long. Bonferroni step-down correction was applied on two-sided hypergeometric test to determine statistical significance. Unless otherwise indicated, all other statistical analyses were performed using GraphPad Prism (v9.1.0) for Windows (GraphPad Software, San Diego, California USA). Data are represented as mean ± standard error of the mean unless otherwise stated.

## Data and Code Availability

RNA-seq, LCM sequencing, and Tuba-seq data are available through Gene Expression Omnibus (GEO) accession number GSE198637. Gene dependency data from the Cancer Dependency Map are publicly available at www.depmap.org. Protein data from UniProtKB are publicly available at www.uniprot.org. All other data are available in the Supplementary Information, or from the corresponding author upon reasonable request.

## ACKNOWLEDGMENTS

We thank Alyssa Ray for administrative support; Dr. Trudy Oliver for sharing the *RPM* mouse model; Pauline Chu and the Animal Histology Service Center at Stanford University for help with histology; the Stanford Shared FACS Facility for flow cytometry services (NIH S10 Shared Instrument Grant S10RR027431-01); the Stanford Veterinary Service Center for expert animal care; the Stanford Genomics Service Center as well as the Protein and Nucleic Acid Facility for help with Bioanalyzer runs; Hyoeun Jung for helping generate illustrations and organize the figures; and all the members of the laboratory of J.S. and M.M.W. for their help and support throughout this study. This work was supported by the NIH (J.S., CA231997 and CA217450; M.M.W. and D.A.P., R01-CA207133, R01-CA231253, R01-CA234349; and J.H.K., F31CA257169-01), the Stanford Cancer Institute (NIH P30-CA124435), and the Agency for Science, Technology and Research (A*STAR) Singapore (Y.T.S.). M.C.L. was supported by Tom and Susan Ford Stanford Graduate Fellowship. H.C. was supported by a Tobacco-Related Disease Research Program (TRDRP) Postdoctoral Fellowship (28FT-0019). C.W.M. was supported by the NSF Graduate Research Fellowship Program and an Anne T. and Robert M. Bass Stanford Graduate Fellowship. C.L. was the Connie and Bob Lurie Fellow of the Damon Runyon Cancer Research Foundation (DRG-2331). J.S. is the Elaine and John Chambers Professor in Pediatric Cancer. The authors would like to acknowledge the NCI Small Cell Lung Cancer Consortium (U24 CA213274) and the AACR Project GENIE registry for sharing SCLC data; interpretations are the responsibility of study authors.

## AUTHOR CONTRIBUTIONS

M.C.L. and J.S. designed most of the experiments and interpreted the results, with help from M.M.W. M.C.L. performed most of the experiments and analysis. H.C., C.W.M., L.A., and M.M.W. helped design and generate Tuba-seq vectors, prepare Tuba-seq libraries from mouse lung, and helped design Tuba-seq experiments and interpret resulting data. C.L. performed the Tuba-seq analysis under supervision of D.A.P. Y.T.S., A.L.H., J.H.K, and G.L.C. helped perform naphthalene injection, virus instillation, and lung collection. A.H. and A.P.D. helped design, generate vectors for, and run the CyTOF for the epitope-tagged NCI-H82 competition assay. C.K. performed the histopathological analysis. S.Z., C.Z., and J.W. performed the laser capture microdissection experiments under supervision of M.V.D.R. M.C.L. and J.S. wrote the manuscript with contributions from all authors.

## COMPETING INTEREST DECLARATION

The authors declare no competing interests.

---

**Supplemental Figure S1:**
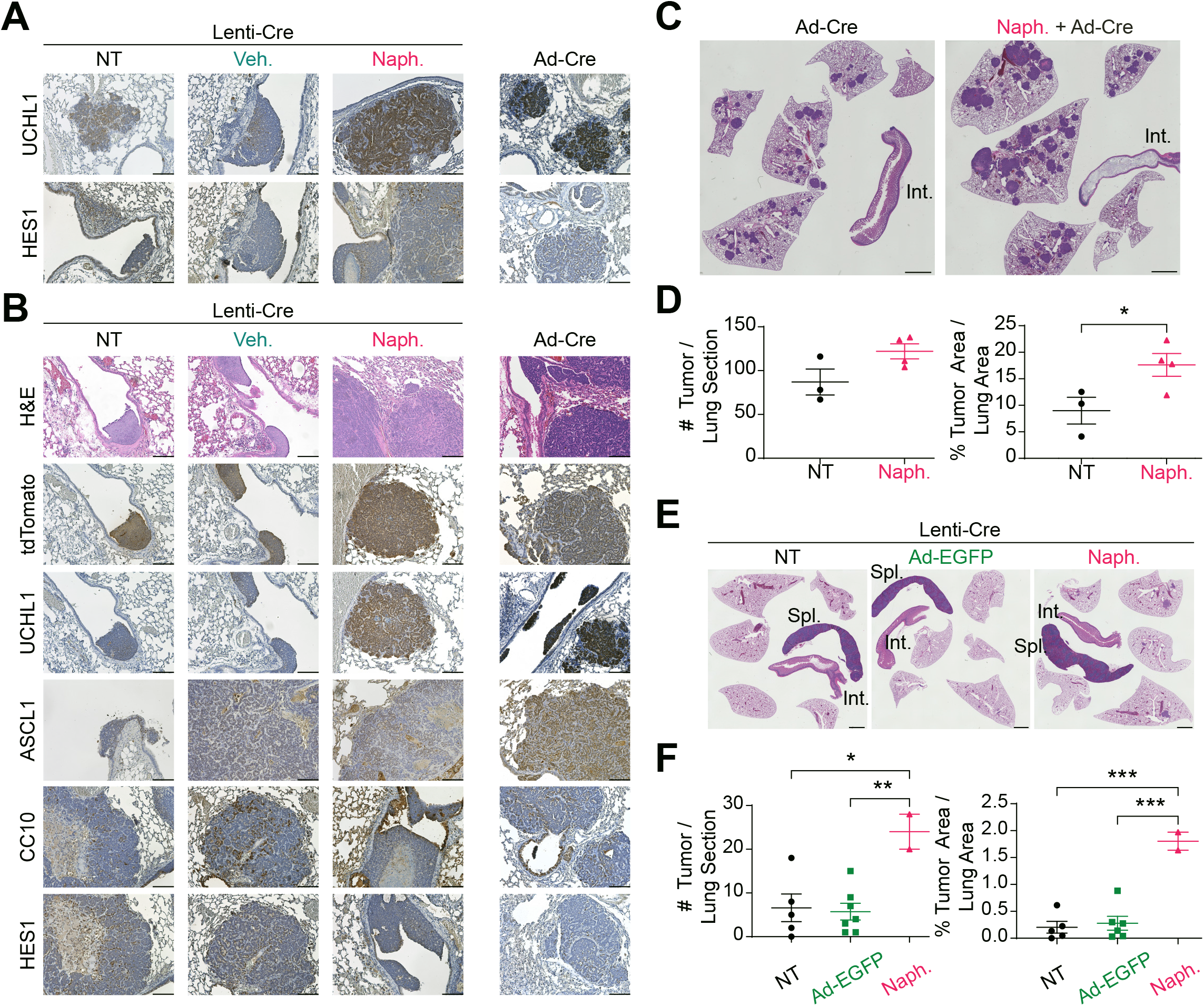
Naphthalene treatment, but not adenoviral inflammation, enhances SCLC development in adenoviral and lentiviral models of SCLC. **A,** Representative images from immunohistochemistry staining (IHC, brown signal) for HES1 and UCHL1 expression on lung sections from mice transduced with Ad-CMV-Cre (Ad-Cre) or HIV-PGK-Cre (Lenti-Cre) alone (NT) or following corn oil (vehicle, veh.) or following naphthalene (naph.) pre-treatment two days prior. Scale bar, 100 µm. **B,** Representative hematoxylin and eosin (H&E) and IHC staining images of lung sections (with some intestine, Int., and spleen, Spl.) from mice transduced with FIV-CMV-Cre (Lenti-Cre) or Ad-CMV-Cre (Ad-Cre) as a control. (n=1 independent experiment, with n=2-3 mice per group). Scale bar, 100 µm. **C,** Representative H&E staining images of lung sections (with some intestine, Int.) from mice transduced with Ad-CMV-Cre (Ad-Cre) alone or following naphthalene pre-treatment (naph.). Mice were collected at 20 weeks following intratracheal instillation. Scale bar, 2 mm. **D,** Quantification of results in (C) (n=1 independent experiment, n=3-4 mice). **E,** Representative H&E staining images of lung sections from mice transduced with FIV-CMV-Cre (Lenti-Cre) alone (NT) or following Ad-CMV-EGFP (Ad-EGFP) instillation or naphthalene pre-treatment (naph.) two days prior. Scale bar, 2 mm. Mice were collected at 24 weeks following intratracheal instillation. **F,** Quantification of results in (E) (n=1 independent experiment, n=2-7 mice per group). Statistical significance for (D) and (F) was determined by one-way ANOVA with post-hoc Tukey test (D) and two-sided unpaired t-test (F). *, p<0.05, **, p<0.01, ***, p<0.001. All data represented as mean ± s.e.m.

**Supplemental Figure S2:**
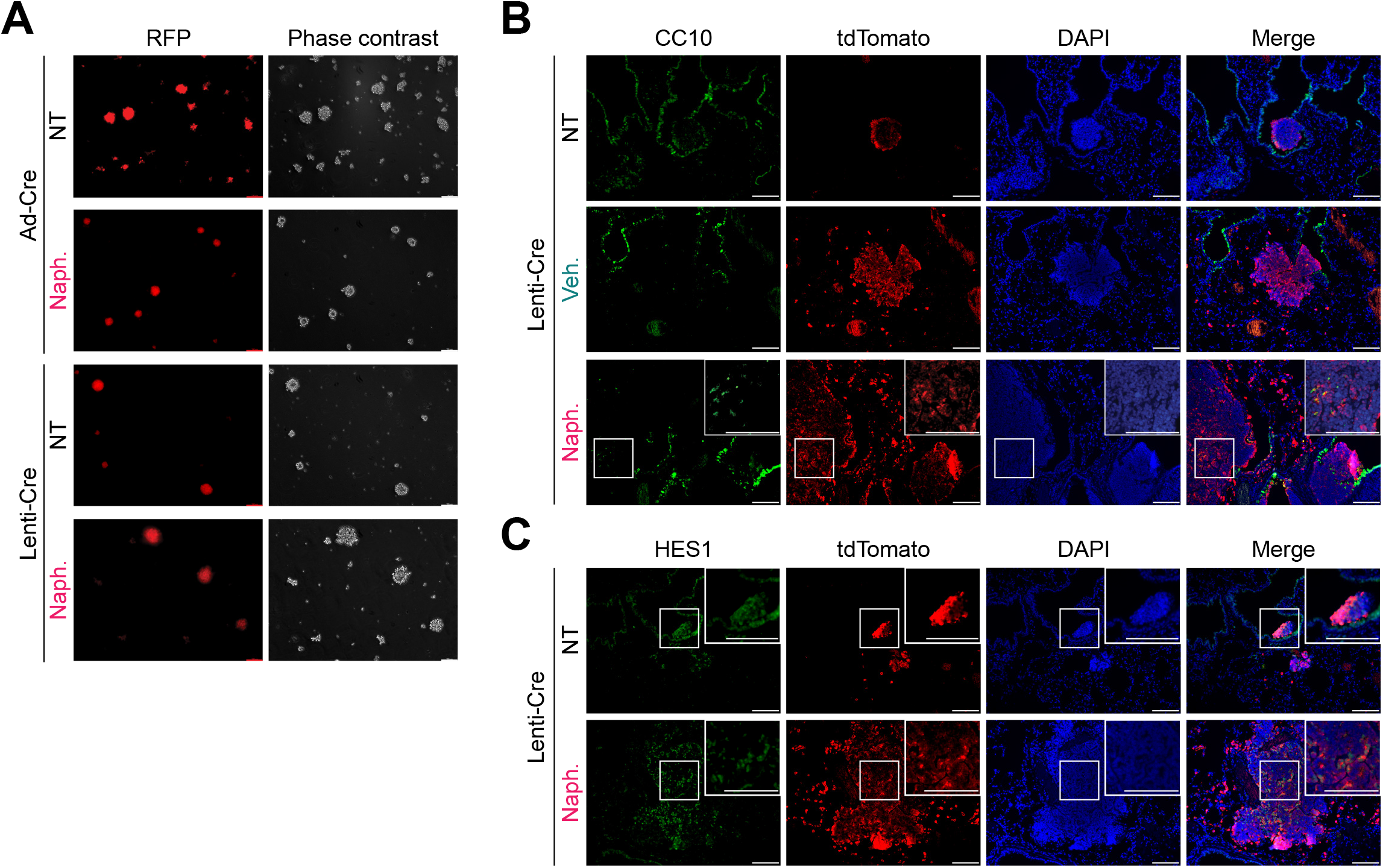
Lenti-Cre-derived *RPR2T* tumors and tumor-derived cell lines show histological and morphological characteristics of SCLC. **A,** Phase-contrast and red fluorescence images of representative *RPR2T* mouse tumor-derived cell lines arising from Adeno-CMV-Cre (Ad-Cre) or HIV-PGK-Cre (Lenti-Cre) transduction alone (NT) or following naphthalene treatment (naph.). Note the growth of spheroids common in neuroendocrine SCLC tumor cell lines. Scale bar, 100 µm. **B,** Representative immunofluorescence images of lung sections from *RPR2T* mice transduced with HIV-PGK-Cre (Lenti-Cre) alone (NT), following corn oil (vehicle, veh.) or following naphthalene pre-treatment (naph.) two days prior. DAPI stains DNA in blue. Scale bar, 100 µm. **C,** Representative immunofluorescence images of lung sections from *RPR2T* mice transduced with HIV-PGK-Cre (Lenti-Cre) alone (NT) or following naphthalene pre-treatment (naph.) two days prior. DAPI stains DNA in blue. Note that CC10^high^ and HES1^high^ cells (in B and C, respectively) within tumors co-stain with tdTomato, indicating that these non-neuroendocrine cells in the Lenti-Cre tumors represent *bona fide* SCLC cells rather than infiltrating, non-transformed cells. Scale bar, 100 µm.

**Supplemental Figure S3:**
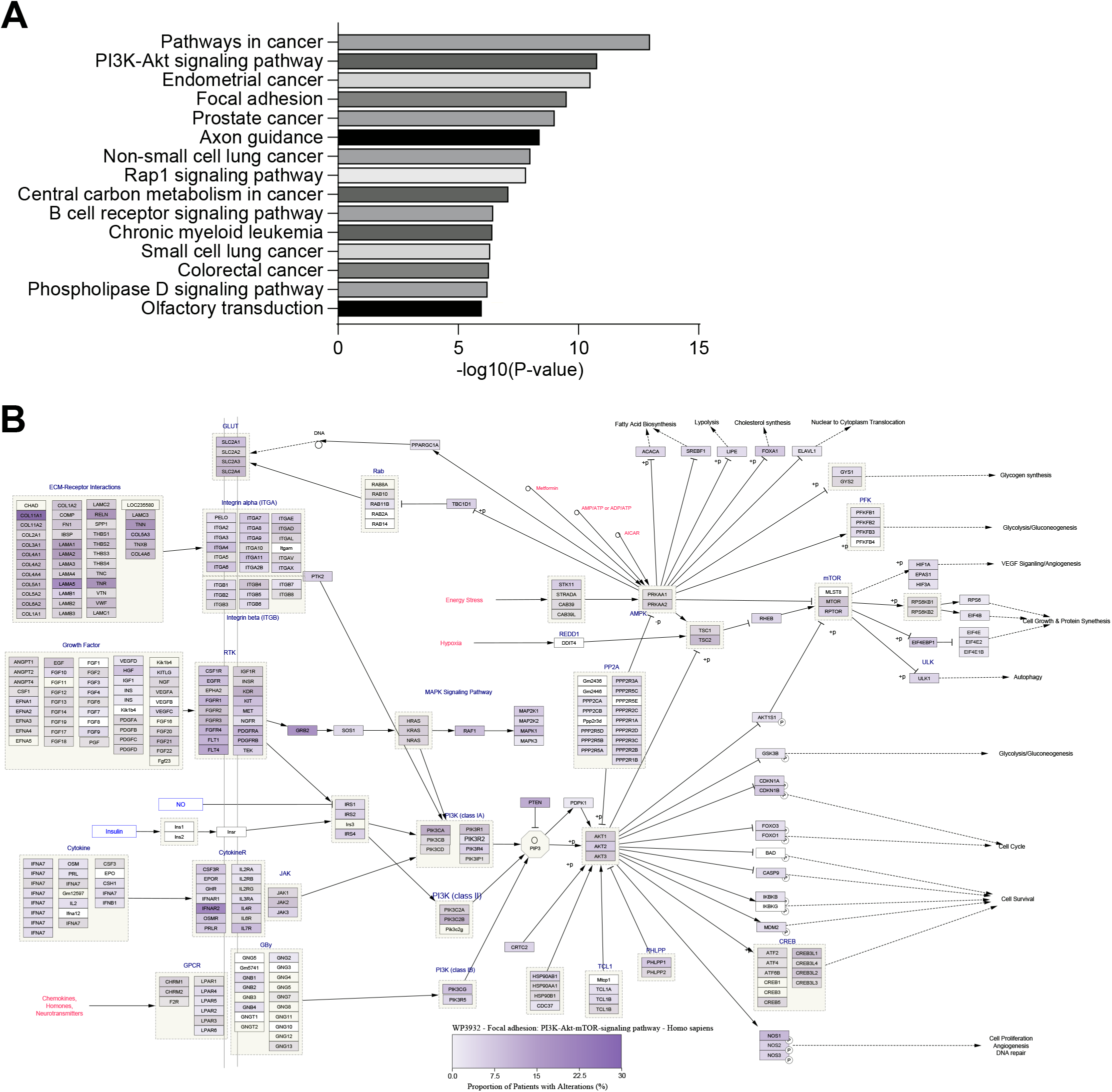
A meta-analysis of SCLC literature reveals enrichment of Focal Adhesion-PI3K-Akt-mTOR pathway member alterations. **A,** Top 10 enriched KEGG pathways for genes altered in ≥3% of SCLC patients, were profiled in at least 250 patients, and coded for protein with amino acid residue length of ≤2,000. Changing the gene cut-off criteria (e.g. removing amino acid residue length limits on protein products and keeping only genes that are expressed at ≥5 RPKM in human SCLC) did not strongly impact the pathway enrichments. **B,** Diagram of Focal Adhesion-PI3K-Akt-mTOR pathway member alteration rates. Pathway network was obtained from WikiPathways using Cytoscape. Fill color indicates % of patients with alterations in that gene. p-value was determined through Bonferroni step-down correction on two-sided hypergeometric test (A).

**Supplemental Figure S4:**
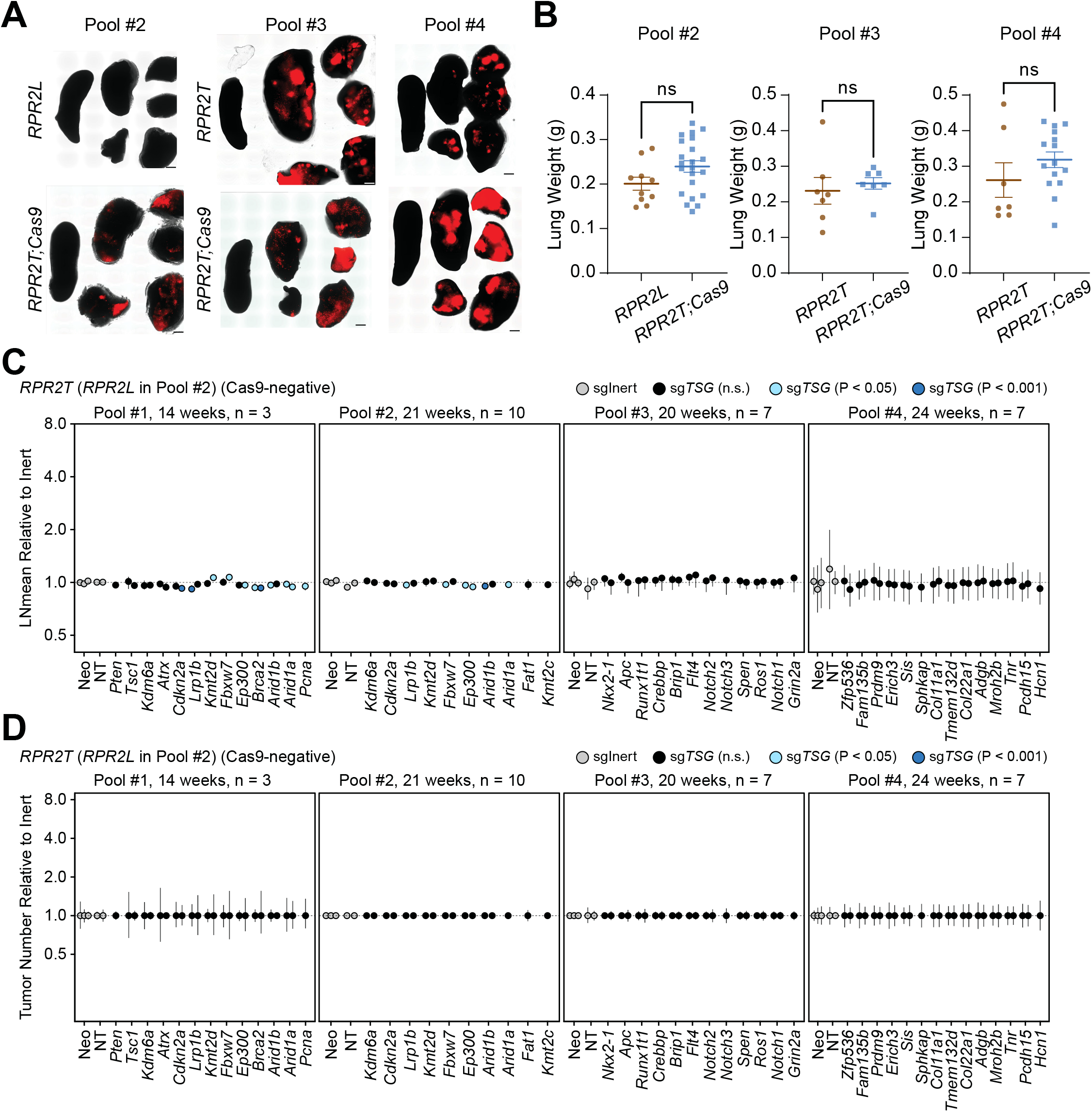
*In vivo* CRISPR screen reveals no sgRNA-dependent enrichment or depletion in *RPR2T* mice lacking Cas9. **A**, Representative tdTomato fluorescence images of tissues isolated from mice transduced with Pools #2, 3, 4. *RPR2;R26^LSL-Luciferase^* (*RPR2L*) mice was used as a control instead of *RPR2T* mice in Pool #2 due to availability, so these mice appear negative for tdTomato fluorescence. Spleen is shown on the left side of the section as a fluorescence negative control. Scale bar, 2 mm. **B**, Lung weights of mice transduced with Pool #2, 3, and 4 at the time of collection. **C,** Log-normal mean tumor size (normalized to tumors with sgInerts) for each putative tumor suppressor gene targeting sgRNA in *RPR2T* mice. For each gene, each circle represents a unique sgRNA. P-values are indicated with a color code. **D,** Tumor numbers (normalized to tumors with sgInerts as well as tumors in *RPR2T* mice for Pools #1, 3-4 (or *RPR2L* mice for Pool #2) for each putative tumor suppressor gene targeting sgRNA in *RPR2T* mice. For each gene, each circle represents a unique sgRNA. P-values are indicated with a color code. The 95% confidence intervals were calculated by bootstrapping (C, D). P-values were determined through two-sided unpaired t-test (B) or bootstrapping followed by Benjamini-Hochberg correction (C, D). Data represented as mean ± s.e.m. (B) or mean ± 95% confidence interval (C-D). ns: not significant.

**Supplemental Figure S5:**
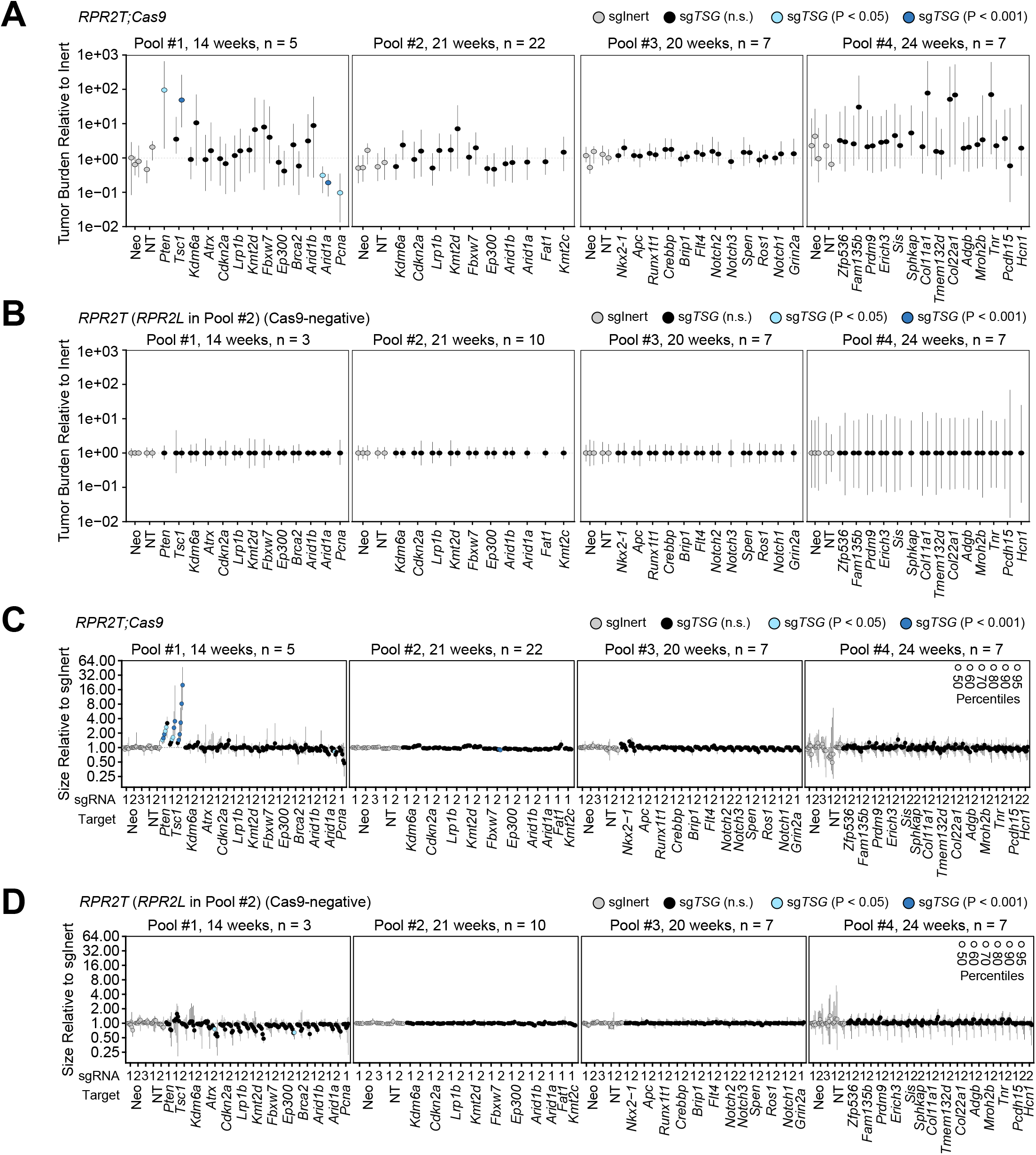
*In vivo* CRISPR screen uncovers both positive and negative effects of gene inactivation on SCLC tumor burden and size in *RPR2* mutant mice. **A-B**, Tumor burden (normalized to tumors with sgInerts and to tumors in *RPR2T* mice) for each putative tumor suppressor gene targeting sgRNA in *RPR2T;Cas9* (A) and *RPR2T* (B) mice. For each gene, each circle represents a unique sgRNA. P-values are indicated with a color code. **C-D,** Tumor sizes at the indicated percentiles (normalized to tumors with sgInerts) for each putative tumor suppressor gene targeting sgRNA in *RPR2T;Cas9* (C) and *RPR2T* (D) mice. For each gene, each circle represents a unique sgRNA. P-values are indicated with a color code. The 95% confidence intervals were calculated by bootstrapping. All p-values were determined through bootstrapping followed by Benjamini-Hochberg correction. Data represented as mean ± 95% confidence interval (A-D).

**Supplemental Figure S6:**
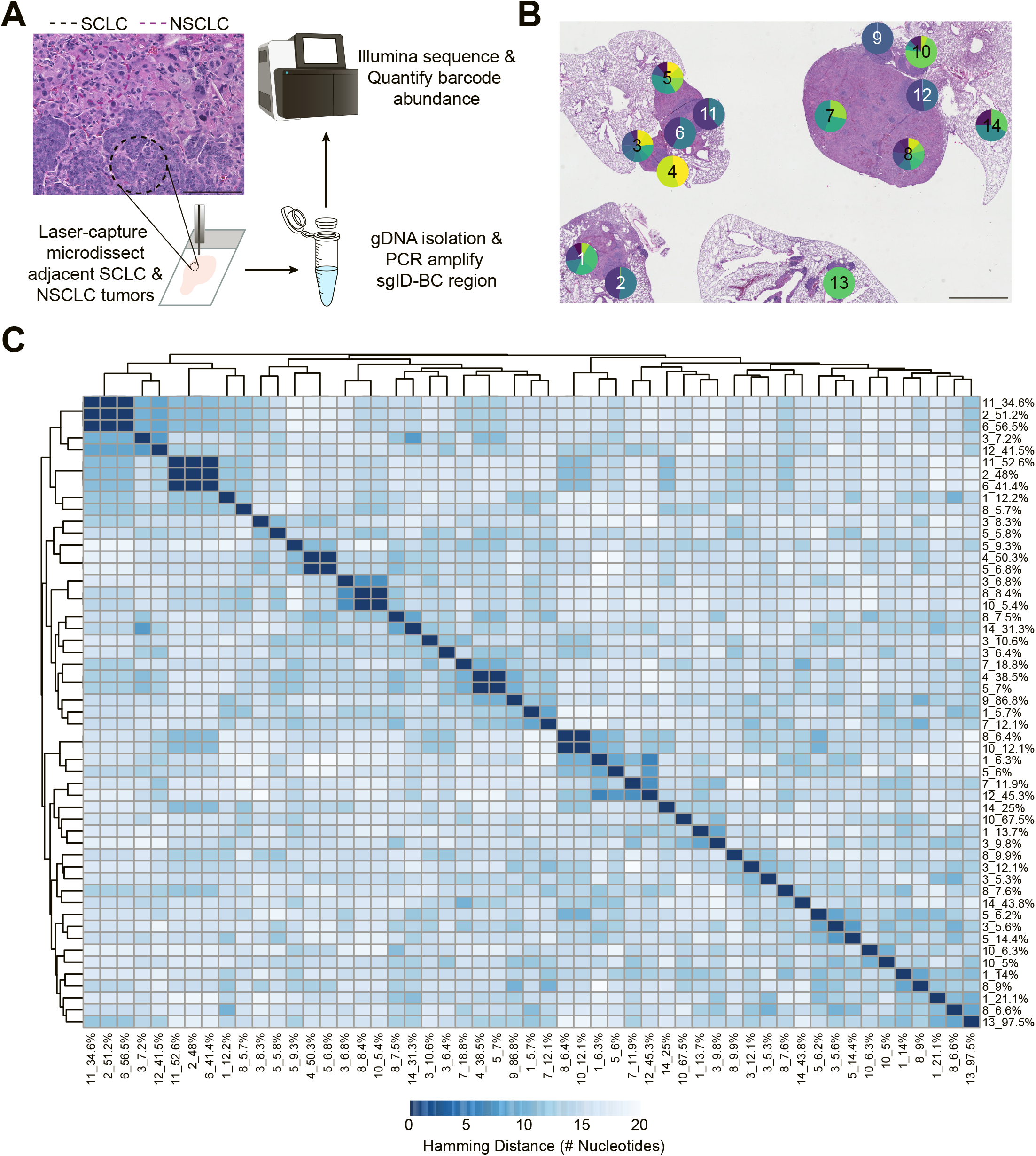
SCLC and NSCLC tumors arising from *Tsc1* inactivation are not clonally related. **A,** Schematic of the workflow for barcode sequencing of laser-capture microdissected tumor sections, with a representative H&E image of SCLC/NSCLC tumors bordering one another. Scale bar, 1 mm. **B,** Hematoxylin and eosin (H&E) staining of the lung used for laser-capture microdissection (n = 1, *RPR2T;Cas9* mice transduced with Lenti-sg*Tsc1*-2/Cre). Section label and representative pie plot of major barcode representation (comprising >5% of barcode frequency from that section) are overlaid on the microdissected area. Scale bar, 2 mm. **C,** An unsupervised clustering of hamming distance between major barcodes (comprising >5% of barcode frequency from that section) in laser-capture microdissected samples. Label indicates sample number and the percent representation of the given barcode in the sample. A hamming distance of 0 indicates an identical barcode.

**Supplemental Figure S7:**
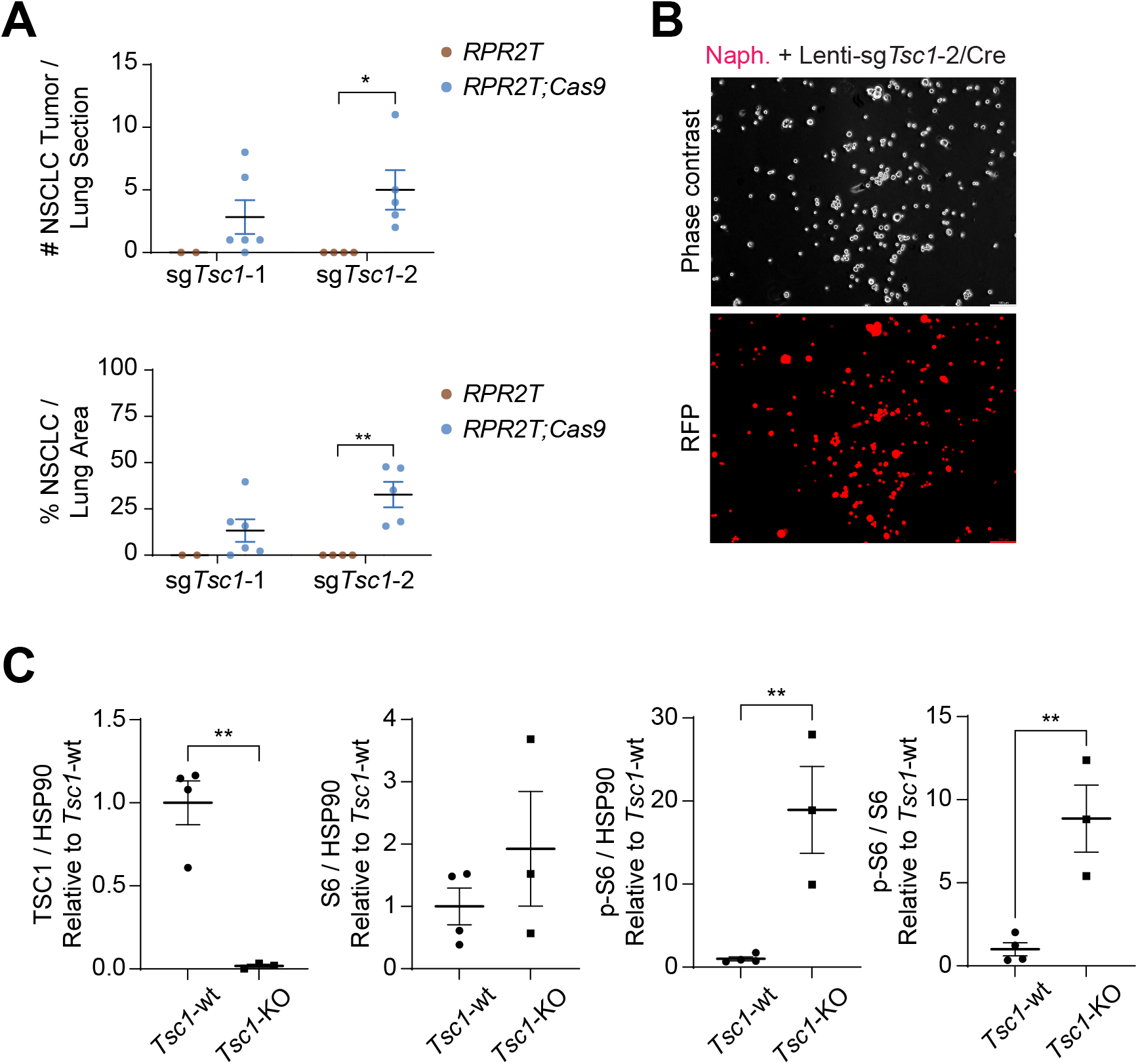
*Tsc1* inactivation generates both SCLC and NSCLC tumors with increased levels of S6 phosphorylation. **A,** Quantification of NSCLC tumor size and number in Figure 4A. **B,** Phase-contrast and red fluorescence images of representative *RPR2T;Cas9* mouse tumor-derived cell lines arising from Lenti-sg*Tsc1*/Cre guide #2 (Lenti-sg*Tsc1*-2/Cre) transduction following naphthalene treatment (naph.) (n = 3). Scale bar, 100 µm. **C,** Quantification of TSC1, S6, and phosphorylated S6 (p-S6) expression from 4D. Values were normalized to *Tsc1*-wt cell lines. Data represented as mean ± s.e.m. for n=2-6 mice (A) or n=3-4 cell lines derived from independent tumors (C) per condition. All p-values were determined through two-sided unpaired t-test. *, p<0.05, **, p<0.01.

**Supplemental Figure S8:**
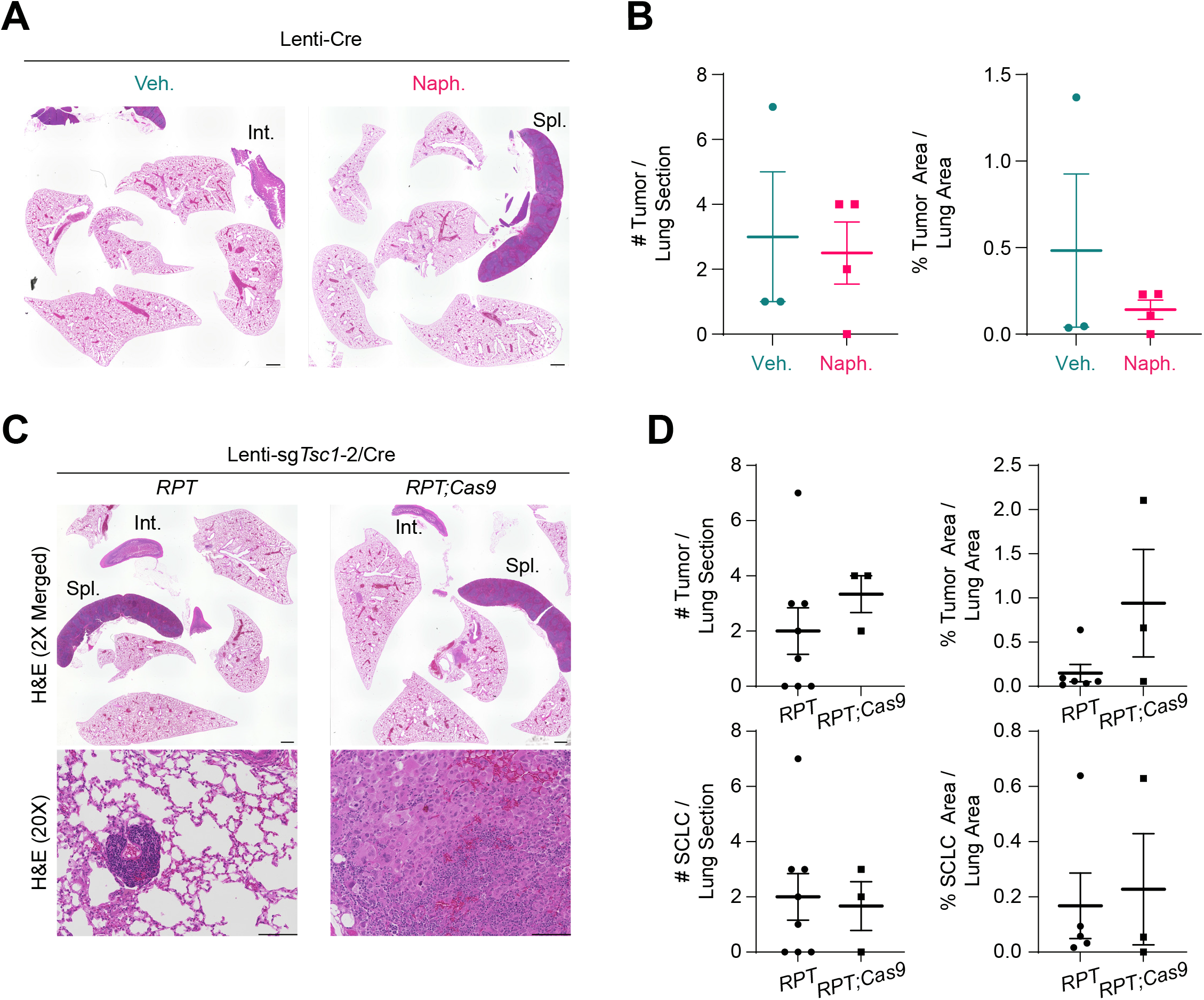
Few tumors are generated in the *RP* mouse model upon transduction with Lenti-Cre. **A,** Representative hematoxylin and eosin (H&E) staining of lung sections (with some intestine, Int., and spleen, Spl.) from *Rb1^fl/fl^*;*Trp53^fl/fl^*;*R26^LSL-tdTomato^*;*H11^LSL-Cas9^* (*RPT;Cas9*) mice at 42 weeks following transduction with HIV-PGK-Cre (Lenti-Cre) post corn oil (vehicle, veh.) or naphthalene (naph.) pre-treatment (n=1 independent experiment, n=3-4 mice per group). Scale bar, 1 mm. **B,** Quantification of tumor burden and numbers from mice in (A). **C,** Representative H&E images from lungs (with spleen and intestine) of *RPT* and *RPT;Cas9* mice transduced with Lenti-sg*Tsc1*/Cre guide #2 (Lenti-sg*Tsc1*-2/Cre) virus (n=1 independent experiment, n=3-8 mice per group). Scale bar, 100 µm. **D,** Quantification of tumor burden and numbers from mice in (C). Data represented as mean ± s.e.m. All p-values were determined through two-sided unpaired t-test.

**Supplemental Figure S9:**
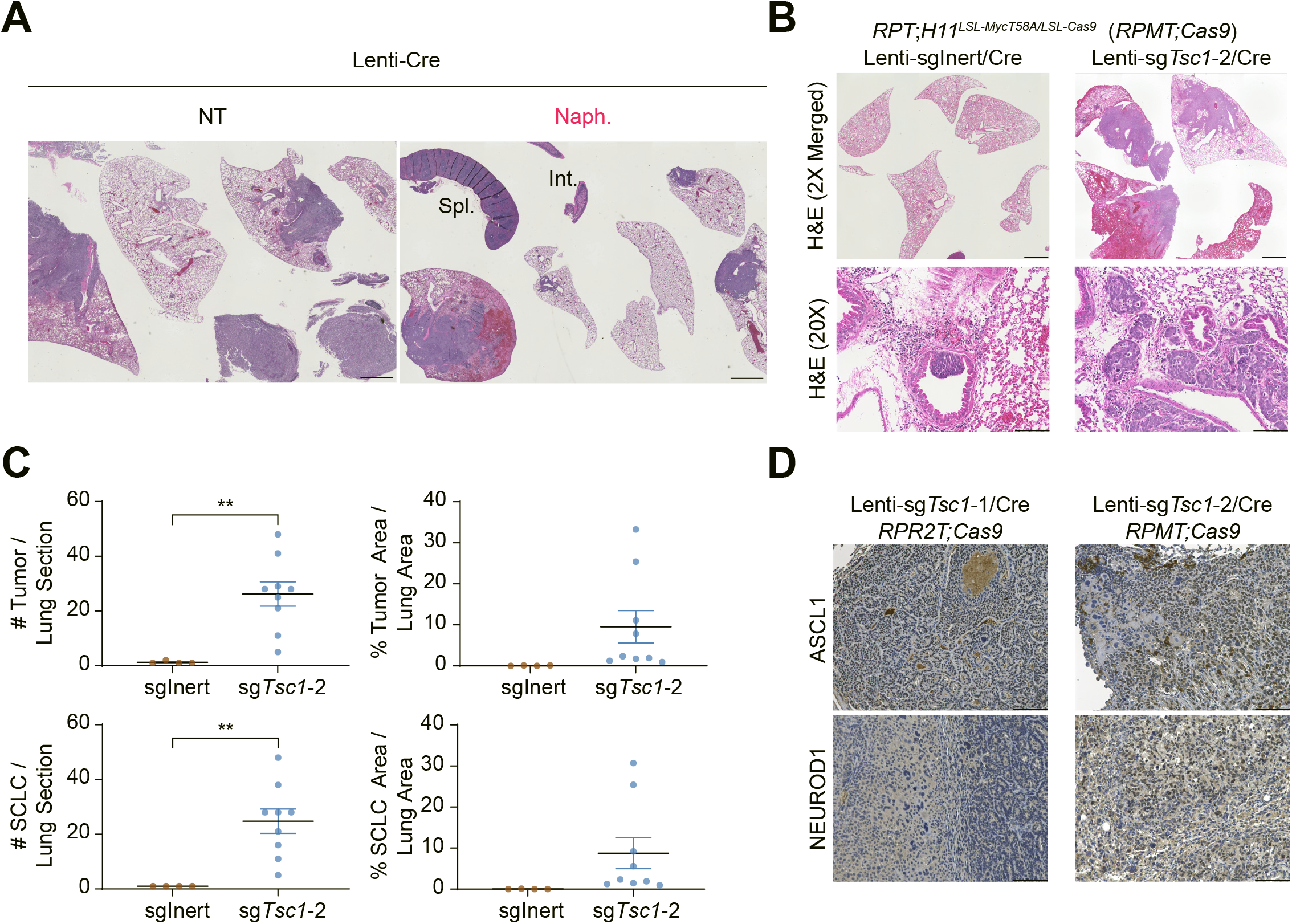
*Tsc1* is a tumor suppressor in SCLC-N in addition to SCLC-A. **A,** Representative H&E images from lungs (with some intestine, Int., and spleen, Spl.) of *Rb1^fl/fl^*;*Trp53^fl/fl^*;*H11^LSL-MycT58A^* (*RPM*) mice transduced with FIV-CMV-Cre (Lenti-Cre) virus without treatment (no treatment, NT) or following naphthalene pre-treatment (naph.) (n=1 experiment, n=2 mice per group). Scale bar, 2 mm. **B,** Representative H&E images from lungs (with spleen and intestine) of *Rb1^fl/fl^*;*Trp53^fl/fl^*;*R26^LSL-tdTomato/+^*;*H11^LSL-MycT58A/LSL-Cas9^* (*RPMT;Cas9*) mice transduced with HIV-sg*Neo1*/Cre (Lenti-sgInert/Cre) or Lenti-sg*Tsc1*/Cre guide #2 (Lenti-sg*Tsc1*-2/Cre) virus (n=1 experiment, with n=4-9 mice per group). Scale bar, 2 mm for 2X Merged panel and 100 µm for 20X section. **C,** Quantification of tumor burden and numbers from mice in (B). **D,** Representative immunohistochemistry staining (IHC, brown signal) images of lung sections from *RPR2T;Cas9* or *RPMT;Cas9* mice transduced with Lenti-sg*Tsc1*/Cre guide #1 (Lenti-sg*Tsc1*-1/Cre) or Lenti-sg*Tsc1*/Cre guide #2 (Lenti-sg*Tsc1*-2/Cre). Scale bar, 100 µm. Data represented as mean ± s.e.m. All p-values were determined through two-sided unpaired t-test. *, p<0.05, **, p<0.01.

**Supplemental Figure S10:**
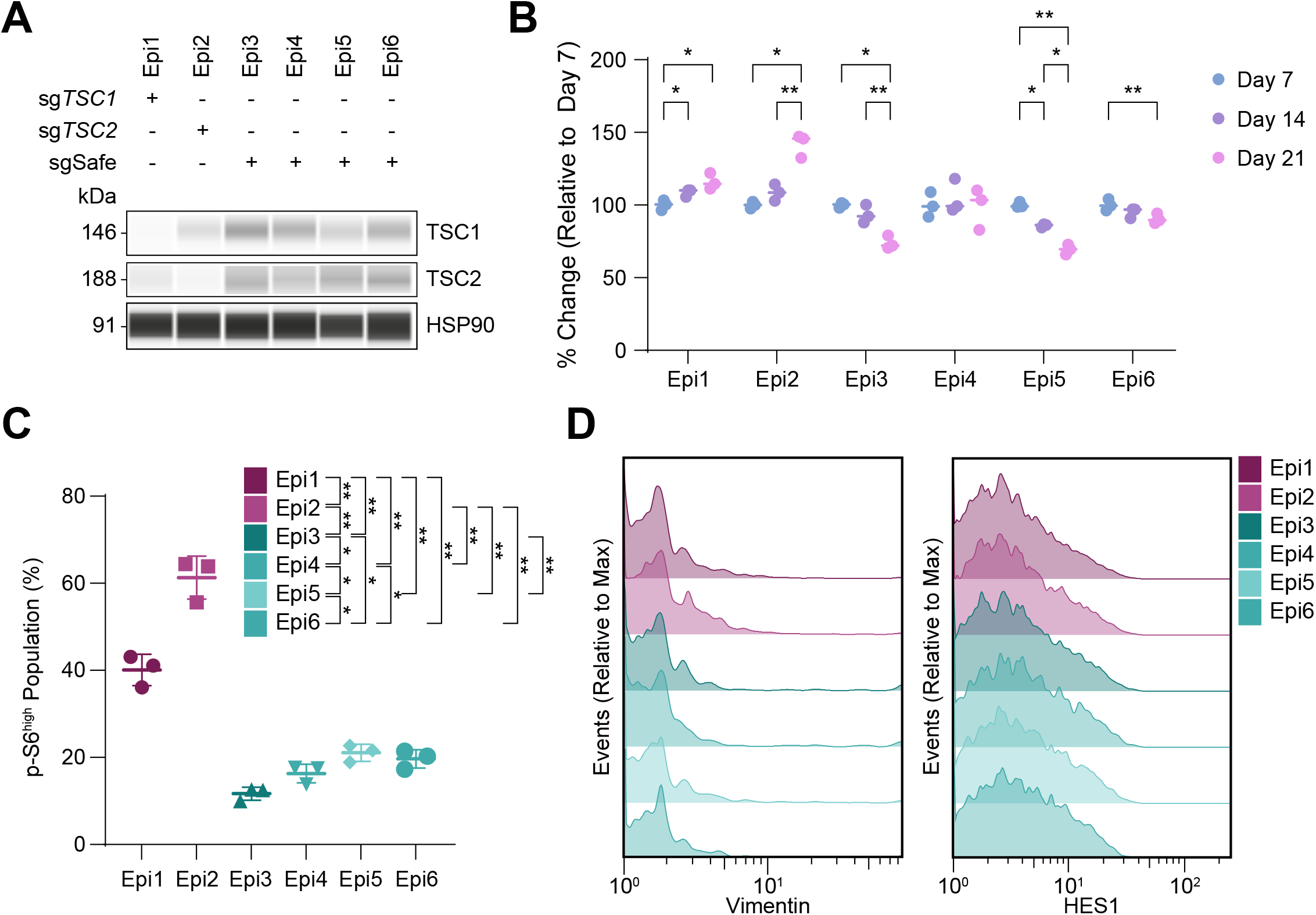
*TSC1*-KO and *TSC2*-KO epitope-combinatorial-tagged NCI-H82 cell lines proliferate faster but do not exhibit non-neuroendocrine differentiation. **A,** Immunoassay of TSC1 and TSC2 in cell lines derived from epitope-tagged H82 cell lines ribonucleofected with indicated sgRNA. HSP90 was used as a loading control. **B,** Representation of data in Figure 4F as percent change in epitope representation within the pool relative to Day 7 (n=1 independent experiment, n=3 technical replicates per day) . **C,** Proportion of p-S6^high^ population in Day 21 samples (n=1 independent experiment, n=3 technical replicates) across epitope-labeled populations. **D,** Modal distribution of vimentin and HES1 signal across the different epitope-labeled populations. One representative experimental replicate from Day 21 is shown; all other replicates across days exhibit similar signal distribution to what is shown. All data represented as mean ± s.d. p-values were determined through repeated measures two-way ANOVA with Geisser-Greenhouse correction followed by post-hoc Tukey test (B-C). *, p<0.05; **, p<0.01; ***, p<0.001.

## Notes

### Competing Interest Statement

The authors have declared no competing interest.

